# De Novo multi-omics pathway analysis (DMPA) designed for prior data independent inference of cell signaling pathways

**DOI:** 10.1101/2022.02.05.479228

**Authors:** Katri Vaparanta, Johannes A. M. Merilahti, Veera K. Ojala, Klaus Elenius

**Author notes:** Address correspondence to: Klaus Elenius.

## Abstract

New tools for cell signaling pathway inference from multi-omics data that are independent of previous knowledge are needed. Here we propose a new de novo method, the de novo multi-omics pathway analysis (DMPA), to model and combine omics data into regulatory complexes and pathways. DMPA was validated with publicly available omics data and was found accurate in discovering protein-protein interactions, kinase substrate phosphosite relationships, transcription factor target gene relationships, metabolic reactions, epigenetic trait associations and signaling pathways. DMPA was benchmarked against existing module and network discovery and multi-omics integration methods and outperformed previous methods in module and signaling pathway discovery especially when applied to datasets with low sample sizes and zero-inflated data. Transcription factor, kinase, subcellular location and function prediction algorithms were devised for transcriptome, phosphoproteome and interactome regulatory complexes and pathways, respectively. To apply DMPA in a biologically relevant context, interactome, phosphoproteome, transcriptome and proteome data were collected from analyses carried out using melanoma cells to address gamma-secretase cleavage-dependent signaling characteristics of the receptor tyrosine kinase TYRO3. The pathways modeled with DMPA reflected both the predicted function and the direction of the predicted function in validation experiments.

## Introduction

Different approaches to model omics data from different levels of cell signaling into cell signaling networks have been developed to understand the interplay, combined effect and causality of cell signaling changes. Previous approaches may be divided into two main categories: methods that are reliant on or independent of previous knowledge. The methods that rely on previous knowledge include the classical gene set enrichment analysis [1], its derivatives, and multiple topology-based methods that use previous data as a backbone to infer pathway networks [2]. Modeling based on previous knowledge poses three major caveats. Firstly, previous knowledge is often incomplete signifying that all present genes, proteins, transcripts and post-translational modification sites are often not covered by the available databases. Secondly, the uneven acquirement of previous knowledge leads to skewed results in endorsing relationships that have already been discovered and extensively researched before. Thirdly, molecular associations are often context-dependent indicating that the associations found in a different context may not universally apply.

Robust methods that are independent on previous knowledge have been generally based on only one measure of molecular association, mainly correlation or partial correlation, between the expression level of different cell signaling molecules [3–6]. Approaches that combine more than one robust molecular association measures such as correlation have not been sufficiently explored. More sophisticated efforts utilizing machine learning methods such as regressions, Bayesian networks and ODE models have been developed to solve single signaling pathway modeling problems such as gene regulatory networks [7,8], but the transferability of these methods to multi-omics data is largely unknown. Most accurate renderings of cell signaling networks have been acquired with methods that are tracking changes in time-course data [7,9]. Unfortunately, the cost and feasibility of acquiring time-course data present a limitation especially in patient samples and in animal models.

The current array of multi-omics integration methods mainly consist of dimensionality reduction and clustering methods that aim to stratify multi-omics data according to disease subtype or patient characteristics and assist in biomarker discovery [10–12]. The module discovery methods such as WGCNA and correlation networks have been expanded and applied to multi-omics data [13,14], but beyond these efforts the multi-omics integration methods that aim to learn cell signaling networks seem to rely on previous knowledge [15–20]. A need for a robust scalable benchmark method that can handle different omics data, is able to track molecular associations from single time point omics data, is independent on previous knowledge and outperforms methods that are based on more than one measure of molecular association, is evident.

## Results

### The underlying concepts of the de novo pathway analysis (DMPA)

We set out to develop a new robust de novo method to infer regulatory modules from different omics data and to combine the modules into multi-omics pathways. Literature search of previous reports of the key features of omics data predicting molecular associations produced the following observations: First, multiple sources indicate the ability of correlation of expression values to predict molecular association to a certain extent [21,22]. Secondly, the findings from an extensive human interactome study indicate that conserved stoichiometry between two proteins inside the same sample can be predictive of protein-protein interaction networks [23]. From the grounds of these two concepts, the basis for the combined score between two proteins, transcripts or phosphosites in an omics dataset was devised to reflect the strength of association. In Figure 1A the mathematical formulation for the correlation and stoichiometry score is presented. The correlation score was based on the nonparametric Spearman correlation which was consequently ranked and adjusted to values between 1 and 0 to allow for a linear scale. The stoichiometry score was based on the relation of the third quartile and first quartile value of relative abundances of the two cell signaling molecules or their modifications in an omics dataset and similarly ranked and adjusted to a linear scale from 1 to 0. The combined score was expressed as a simple non-weighted multiplication of the correlation and stoichiometry scores similarly ranked and adjusted to a linear scale between 1 to 0. Nonparametric formulation was chosen to overset technical variation and to desensitize the model to outlier values.

**Figure 1.**
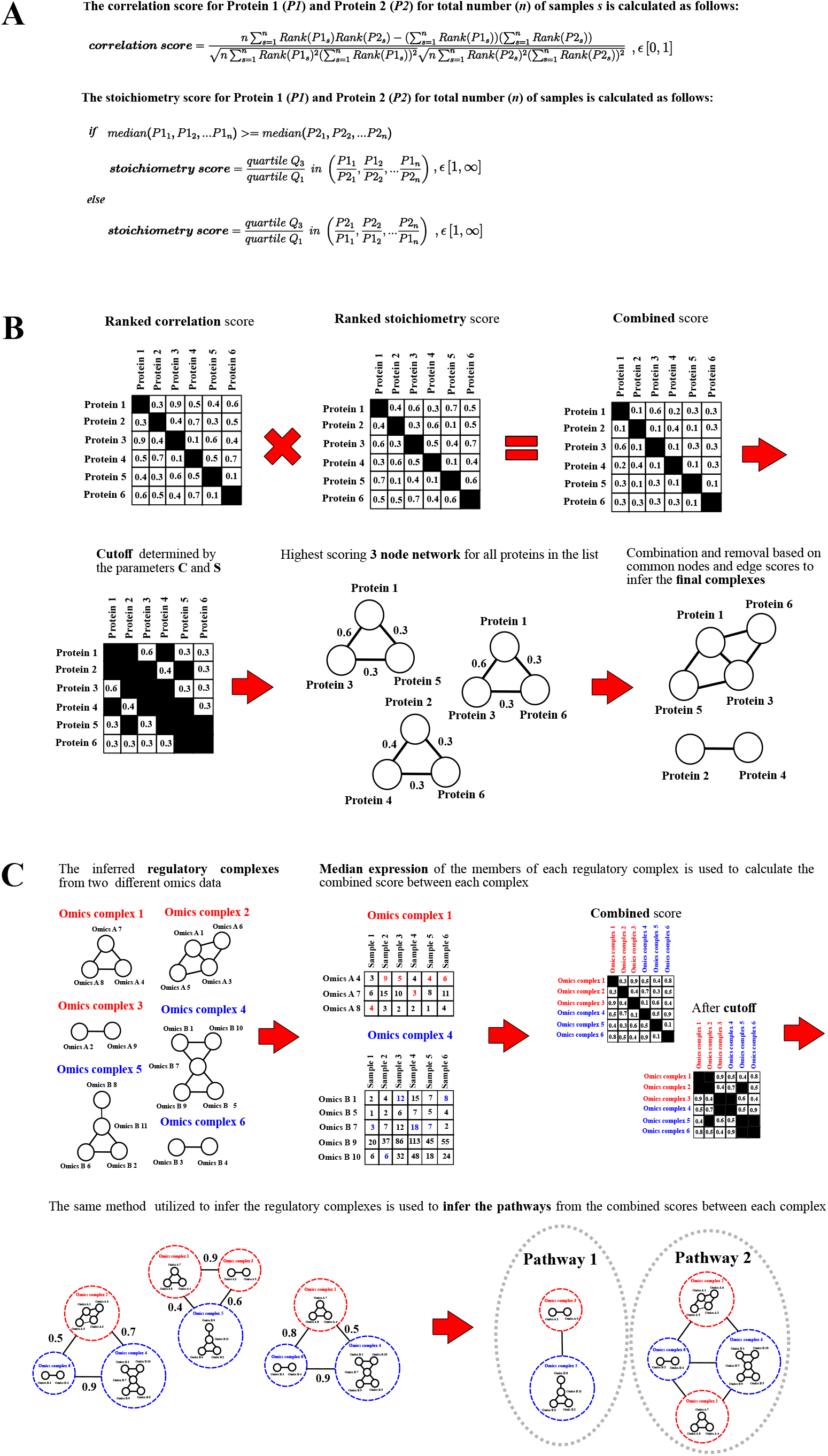
The underlying concepts of the de novo multiomics pathway analysis. **A:** Mathematical formulation of the correlation (Spearman correlation) and stoichiometry scores. **B:** Schematic representation of the inference of regulatory omics complexes in the de novo multi-omics pathway analysis. **C:** Schematic representation of the principles of combining the regulatory omics complexes into pathways.

To reduce computational demand (O(n^2^) for run time), a cut-off for the combined score was devised to consider only strong associations. In the current version of the inference method the cut-off parameter is controlled by two parameters that reflect the number of associations considered for the molecules in the data (parameter C) and the predicted amount of non-associating features in the data (parameter S).

The inference of regulatory modules from the combined score after the cutoff was designed to follow the nearest neighbor principle, which assumes that the combination of combined scores between 3 cell signaling molecules is more predictive than a single combined score between two molecules or their states in the dataset. The inference method is outlined in Figure 1B and utilizes the combined scores above the chosen cut-off threshold to find maximal scoring 3 member cliques. These 3 member cliques are consequently combined to find larger complexes and trimmed to remove members that are present in a higher scoring complex. The principles of the combination and trimming are outlined in more detail in the Materials and methods section.

The combination of the regulatory complexes was designed to follow a similar principle to the initial inference of the regulatory complexes (Figure 1C). A median expression value of all the members of a complex in each sample is used as a representation of the expression values of a complex and the combined score is calculated based on these values. To allow equal combination of down-and up-regulated complexes, an absolute value was taken from the Spearman correlation to equally weigh positive and negative correlations. The complexes were then combined further into pathways based on the combined scores above a certain threshold with the same rules for combining and trimming the maximal scoring 3 complex cliques that were used in the initial regulatory complex inference. The de novo mulri-omics pathway analysis is available both as matlab code and executables in Mendeley Data: https://data.mendeley.com/datasets/m3zggn6xx9/draft?a=71c29dac-714e-497e-8109-5c324ac43ac3.

### Validation of regulatory complexes modeled with DMPA

The performance of the derived regulatory complex inference method was tested on available interactome [23], phosphoproteome [24], and transcriptome [25] data. While molecular associations are context-dependent, certain associations are more frequently reproduced in different contexts and sometimes even stable. Therefore, a level of conservation is expected from a score that would accurately reflect the probability of molecular association. The conservation of the correlation, stoichiometry and combined scores were examined by modeling a set of proteins, phosphorylation sites or transcripts from one dataset into regulatory complexes and examining the median of the corresponding scores derived from the same complexes in another dataset. A median score above 0.5, the median score expected if the scores would be randomly assigned to molecular associations, indicated conservation.

In all examined omics data, the combined score was statistically significantly conserved, while the stoichiometry score was significantly conserved only in the mass spectrometry-derived interactome and phosphoproteome data (Figure 2A). The correlation score in turn was below the 0.5 line in all tested omics data, indicating that the correlation score alone is least likely to predict molecular associations (Figure 2A).

**Figure 2.**
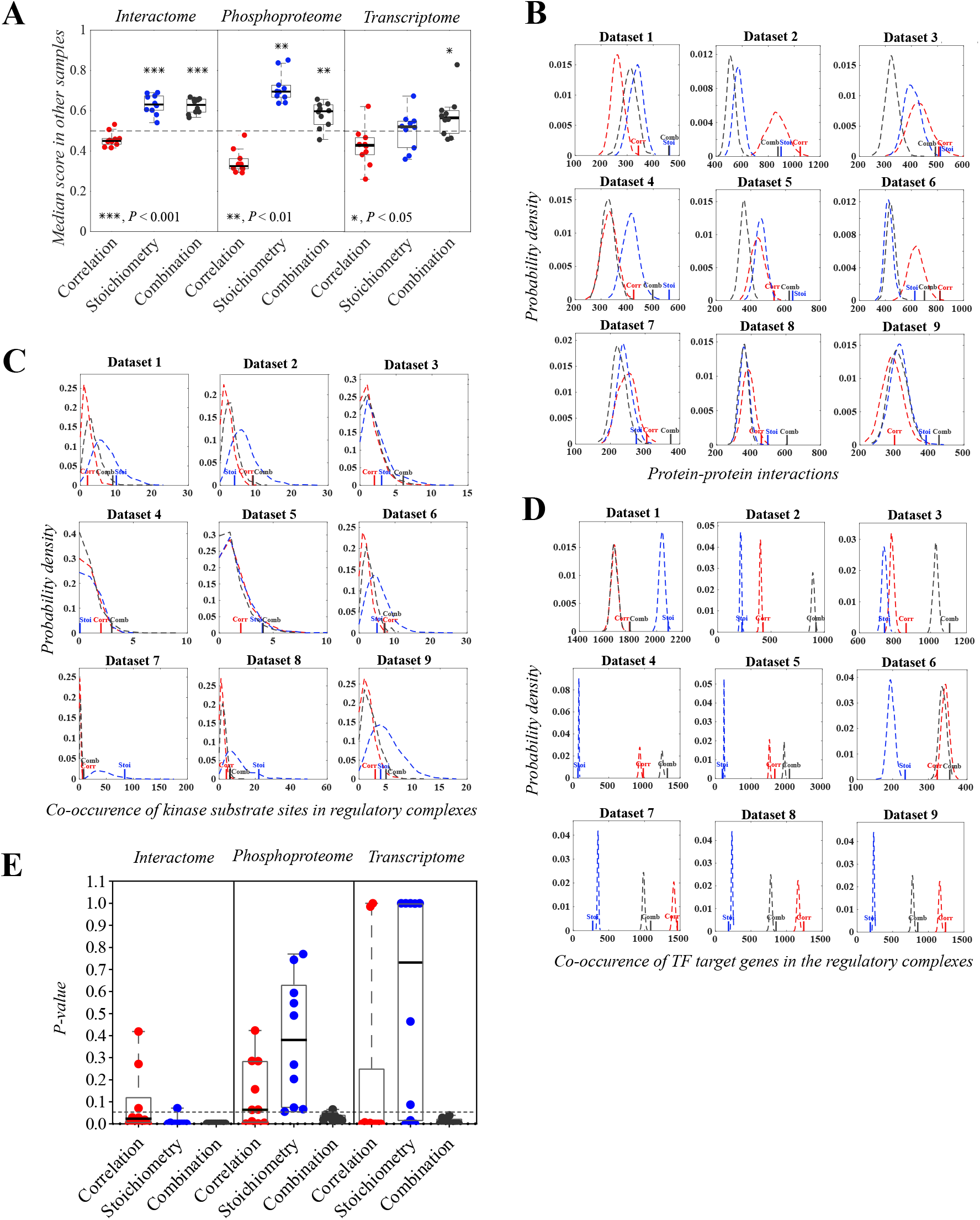
Validation of regulatory complexes modeled with the DMPA. **A:** The conservation of correlation, stoichiometry and combined score in interactome, phosphoproteome and transcriptome data. One dot represents the median score between all complex members of all modeled complexes of a dataset that was not utilized for the initial regulatory complex inference. The dashed line represents the median score of randomized complexes. For statistical testing, two-tailed one sample T-test against the theoretical median value of 0.5 was utilized. **B-E:** Empirical probability densities simulated by randomizing the members of the modeled regulatory complexes into complexes of the same size and the corresponding P-values for the complexes modeled from either interactome (B, E), phosphoproteome (C, E) or transcriptome data (D, E) by either correlation (Corr), stoichiometry (Stoi) or combination score (Comb). The sum of co-occurrences of either interacting proteins as determined by the STRING database, phosphorylated kinase substrate residues as determined by the PhosphoSitePlus database or transcription factor target genes as determined by the ENCODE database in the modeled complexes were used as a validation score. The empirical probability densities are visualized with dashed lines of the corresponding color and the score for the modeled network in solid line. One dot in the boxplot represents one P-value for one dataset and the horizontal line the median value. The datasets were acquired from published omics data. Red: correlation score; blue: stoichiometry score; grey: combined score.

To validate that the regulatory complexes reflect discovered molecular associations, a method to count the number of known associations in the modeled regulatory complexes was devised. Known protein-protein interactions, kinase substrate phosphosite relationships, and transcription factor target gene annotations were downloaded from STRING [26], PhosphoSitePlus [27] and ENCODE [28] databases, respectively. The sums of all co-occurrences of proteins known to interact, substrate sites known to be phosphorylated by the same kinase, and transcripts known to be the target gene for the same transcription factor inside the modeled regulatory complexes were used as validation scores. The validation scores were calculated from the regulatory complexes inferred with correlation, stoichiometry and combined scores to compare the effect of the different scores on the accuracy of the model. The validation scores from the real modeled regulatory complexes were compared to the validation scores of randomly formulated regulatory complexes of the same size to calculate the probability that a similar validation score would be acquired by random. The probability density of the validation scores of these randomized regulatory complexes are indicated with a dashed line and the validation scores for the regulatory complexes modeled with different scores with solid lines in the Figure 2B-D. The corresponding P-values drawn from the cumulative probability densities of the randomized regulatory complexes for the validation scores of the regulatory complexes modeled with different scores are visualized in Figure 2E.

The regulatory complexes modeled with the combined scores consistently reflected known molecular associations in all interactome, phosphoproteome and transcriptome data. In interactome data, the co-occurrence of two proteins in the same regulatory complex predicted a protein-protein interaction between the two proteins more often than random assignment to regulatory complexes of the same size. Additionally, the median STRING score, that reflects the confidence of the interaction based on the amount and type of evidence on the protein-protein interaction, was consistently higher in the regulatory complexes modeled with the combined score than in the randomized regulatory complexes (Supplementary Figure 1). In the phosphoproteome data, the assignment to regulatory complexes modeled with the combined score indicated that the two phosphorylation sites were controlled by the same kinase more often than by random assignment. In the transcriptome data, in turn, two transcripts in the same regulatory complex modeled with the combined score were more likely to be the target gene for the same transcription factor than randomly chosen transcripts. This effect was similarly observed in transcription factor target gene annotations acquired from REMAP and Literature libraries from the ChEA3 database [29] (Supplementary Figure 2). The regulatory complexes modeled with only the correlation or stoichiometry score varied in their ability to find known molecular associations based on the modeled data. The regulatory complexes modeled with stoichiometry score were sufficient, although not as consistent as the regulatory complexes modeled with the combined score, in reflecting known protein-protein interactions in interactome data. However, the regulatory complexes modeled with stoichiometry score failed to reproduce known molecular associations in phosphoproteome and transcriptome data. The regulatory complexes modeled with the correlation score were more or less able to capture known transcription factor target gene relationships in the transcriptome data but were insufficient in finding known protein-protein interactions and kinase substrate relationships in the interactome and phosphoproteome data, respectively. All in all, the regulatory complexes inferred with the combined score consistently captured known molecular associations in all tested omics data, while the complexes inferred with other scores were found lacking in reproducibility and performance across omics datatypes.

To examine the scalability of DMPA to omics data other than the initially tested interactome, phosphoproteome and transcriptome data, the conservation of the combined score was also similarly examined in methylome, metabolomics and protein acetylation data. The combined score was discovered indeed to be significantly conserved also in these omics data (Supplementary Figure 3A). The ability of the regulatory complexes modelled with DMPA to predict known molecular associations from available metabolomics and methylation data was additionally examined. The regulatory complexes modelled with DMPA were able to predict metabolites involved in the same metabolic reaction in metabolomics data and methylated chromatin regions associated with the same trait in methylome data (Supplementary Figure 3B). These data indicate that DMPA is scalable and can additionally be applied to other omics data than the initially validated interactome, phosphproteome and transcriptome data.

### Validation of regulatory complex combination to pathways by DMPA

Since signaling pathways are a combination of molecular associations, they are similarly expected to be partially conserved. The conservation of the regulatory complex combination to pathways was assessed by modeling a multi-omics dataset into regulatory complexes with either correlation, stoichiometry or combination score, combining the regulatory complexes with either correlation, stoichiometry or combined score and examining the median of the corresponding scores derived from the same modeled pathways in another multi-omics dataset. Multi-omics data from normal and cancer tissue samples from the LinkedOmics [30] database were utilized. Of all the tested score combinations for the analysis, inferring the regulatory complexes with the combined score and combining the inferred regulatory complexes into pathways with a combined score was the only score combination for the analysis that demonstrated statistically significant conservation (Figure 3A).

**Figure 3.**
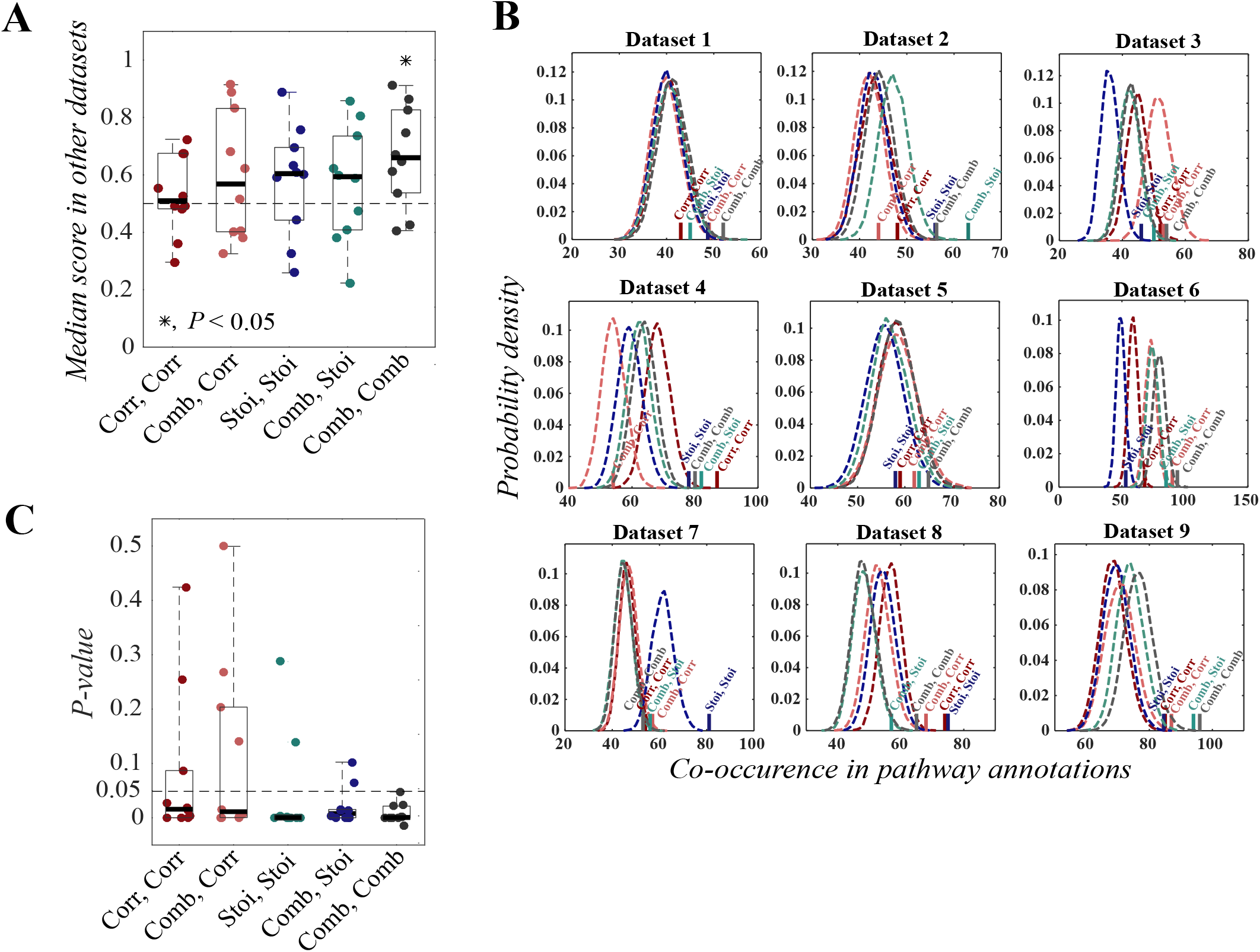
Validation of regulatory complex combination to pathways with DMPA. **A:** The conservation of correlation, stoichiometry and combined score in complexes modeled with either correlation (Corr), stoichiometry (Stoi) or combined (Comb) score and combined with either correlation, stoichiometry or combined score. Corr, Corr: modeled with correlation score, combined with correlation score. Comb, Corr: modeled with correlation score, combined with combined score. Stoi, Stoi: modeled with stoichiometry score, combined with stoichiometry score. Comb, Stoi: modeled with stoichiometry score, combined with combined score. Comb, Comb: modeled with combined score, combined with combined score. One dot in the boxplot represents the median score between all complexes of all inferred pathways of a dataset that was not initially utilized for the pathway inference. The horizontal line represents the median value and the dashed line the median score for randomized pathways. For statistical testing, two-tailed one sample T-test against the theoretical median value 0.5 was utilized. **B-C:** Empirical probability densities simulated by randomizing the modeled regulatory complexes into pathways of the same size and the corresponding P-values. Acetylation, methylation, proteomics, phosphoproteomics, transcriptomics and protein array data from LinkedOmics was modeled and combined. The sum of co-occurrences of genes, transcripts and proteins in a pathway annotation in separate combined regulatory complexes as determined by the PathwayCommons database were used as a validation score. One dot in the boxplot represents a P-value for one dataset and the line the median value. The datasets were acquired from published multi-omics data.

To ensure that DMPA is also able to combine the inferred regulatory complexes into pathways that reproduce known signaling pathways, a validation method was devised. The sum of all co-occurrences of signaling molecules in separate regulatory complexes combined by the DMPA in pathway annotations was used as a validation score for the modeled pathways. The reference pathway annotations were downloaded from the PathwayCommons database [31]. The validation scores were compared to the validation scores calculated from 10,000 randomly combined pathways of the same size to estimate the probability that random assignment of the regulatory complexes would produce pathways that equally reflect known pathway annotations. The probability density of the validation scores of these randomly combined pathways are indicated with a dashed curve and the validation scores for the regulatory complexes modeled with different scores with solid lines in Figure 3B. The corresponding P-values drawn from the cumulative probability densities of the randomized pathways for the validation scores of the regulatory complexes modeled with different scores and combined with different scores are visualized in Figure 3C. The pathways inferred and combined with the combined score consistently reproduced associations represented in known pathway annotations. The second-best performers were the versions of the analysis method where the pathways were inferred with either the stoichiometry score or the combined score and combined with the stoichiometry score. The correlation score-based versions of the analysis method performed worst in finding molecular associations present in known pathway annotations. Taken together, the results from the validations and the level of conservation of the combined score in other datasets strongly imply that the DMPA can find molecular associations and pathways that reflect previously observed cell signaling modules.

### Validation of the design decisions of DMPA

To explore the validity of the design decision of DMPA, the effect of different design decisions of DMPA on the ability of the inferred molecular associations to reflect known molecular associations was examined as in Figure 2B-E. Since the P-values in many cases approached zero the relative distance of the validation score from the median value of the random distribution was used as a measure instead of the P-value like in Figure 2B-E. First, the effect of the choice to rank and adjust the stoichiometry and correlation score on the amount of known molecular association in the inferred molecular associations was assessed. In both interactome and transcriptome data the choice to rank and adjust the stoichiometry and correlation score (combination, ranked) increased the number of known associations in the inferred molecular associations against the version of the combination score where the unadjusted correlation score and the inverse of the unadjusted stoichiometry score were multiplied (combination, raw) (Supplementary Figure 4A). In phosphoproteome data the choice to rank and adjust the correlation and stoichiometry had no significant effect. Second, the effect of the design choice to find the maximally scoring 3 node cliques for each signaling event in the data instead of a linear approach, where for each signaling event only the maximally scoring signaling event is selected was examined. For both interactome and phosphoproteome data searching the maximally scoring 3 node cliques instead of the single maximally scoring signaling event, increased the amount of previously discovered molecular associations in the inferred molecular associations (Supplementary Figure 4B). For transcriptome data the design choice had no significant effect. Taken together the results imply that the design decisions of DMPA have mostly a positive effect or a neutral effect depending on the examined omics data on discovering true positives in the inferred molecular associations. In combination the effect of adhering to both design decisions increased the number of known associations in the inferred molecular associations for all tested omics data types.

### Sensitivity analysis of the parameters of DMPA

To assess the sensitivity of DMPA to parameter choice a sensitivity analyses were conducted. First the effect of parameter choice of the parameters that control the size of the regulatory modules was examined with available interactome data. The change in these parameters affected the size and amount of the regulatory modules but had no negative effect on the number of known associations in the inferred molecular associations against randomly modelled molecular associations (Supplementary Figure 5). This suggests that the parameters can be safely adjusted if larger or smaller modules are desired from the inference method.

Second, the effect of the parameter choice of the parameters C and S that control the cut-off value of the combined score was assessed in both available omics and simulated data (Figure 1B). To examine the effect of the parameter S choice on the amount of known molecular associations in the inferred molecular associations and on the algorithm performance, published transcriptome data was analyzed with different S parameter values. Different S parameter values had a significant effect on the run time but no adverse effect on the number of known molecular associations in the inferred molecular associations (Supplementary Figure 6). To test the effect of the C and S parameters for the true positive and false positive rate DMPA was applied to normally distributed, negative binomially distributed and beta distributed simulated data (Supplementary Figure 7-9). DMPA was able to accurately discover the true modules among the randomly modelled data. The performance of the inference method was, however, discovered to be sensitive to the parameter S to an extent. When the parameter S was set too low the likelihood of false positives in the regulatory modules was increased. In turn when the parameter S was set too high true positives were lost. It is note of that DMPA never misassigned molecules that belong to a regulatory module into another regulatory module, but rather the false positives always came from the simulated pool that had no true module assignment. The proportion of the molecules or their states that have no association with other molecules or their states in the dataset in real omics datasets is expected to be low. The raw correlation and stoichiometry scores of the molecules that were assigned or not assigned into regulatory modules were examined and a significant drop in the maximum raw correlation and inverted raw stoichiometry score was discovered for the molecules that had no true module assignment (Supplementary Figure 10A). To this end a script to suggest values for the parameter S was devised to help the user to set the parameter S (Supplementary Figure 10B). The effect of the parameter C was also examined and it was discovered that the value for parameter C should be set only based on the size of the dataset (3 for small datasets, 30-300 molecules in the dataset; 1 for larger datasets, 500< molecules in the dataset).

Third, the sensitivity of DMPA to the parameter that controls the zero-inflated version of the analysis was assessed. Zero-inflated interactome data and non-zero-inflated transcriptome data was analyzed both with the zero-inflated (weighted stoichiometry score) and non-zero-inflated (non-weighted stoichiometry score) version of DMPA and the amount on known associations in the inferred molecular associations against randomly modelled molecular associations was estimated. In the zero-inflated interactome data (over 30% missing values), the zero-inflated version of the analysis significantly increased the number of known associations in the inferred modules (Supplementary Figure 11). In contrast, in the non-zero-inflated transcriptome data (less than 25% missing values) the zero-inflated version discovered fewer known associations than the non-zero-inflated version.

The choice of zero-inflated and non-zero-inflated version of the analysis has a significant effect on the accuracy of the inference and should be set based on the number of missing values in the dataset.

### DMPA outperforms previous approaches

To assess the performance of DMPA, DMPA was benchmarked against relevant module discovery and multi-omics integration methods. Transcriptome datasets were chosen for the benchmarking of module discovery methods since it allowed the comparison against regulatory network inference methods that are specifically designed to be used with gene expression data. WGCNA [3], LOPC [5] and the DREAM4 challenge winning regulatory network inference method GENIE3 [32] were selected for comparison. Since LOPC returns network edges instead of modules, a separate validation strategy was devised to benchmark against LOPC which is detailed in the Materials and Methods section. For GENIE3 a module was defined as the genes predicted to be controlled by the same upstream gene. Although all WGCNA, LOPC and GENIE3 were able to discover significantly more known molecular associations in the inferred molecular associations than random modelling, DMPA was the most consistent performer across datasets and in most cases discovered most known molecular associations (Figure 4A-B). Strikingly, a clear difference in the performance of DMPA and the other tested methods could be seen with datasets with low sample sizes. To further assess the performance of DMPA and the other module discovery methods with low sample sizes, the module discovery methods were run with datasets with different sample sizes and the amount of known molecular associations in the inferred molecular associations against the randomly assigned molecular associations were examined. DMPA required only 6 samples (3 control and 3 treatment samples) to discover known molecular associations while WGCNA and GENIE3 required over 30 (Figure 4C). An attempt to run zero-inflated interactome data with WGCNA and LOPC was made, but the current R implementations of these approaches were unable to handle the zero-inflated interactome data. DMPA in contrast has the advantage of performing without the imputation of the missing values of the zero-inflated data. These results imply that DMPA is more scalable, has less limitations and can discover more known associations than the tested module and network discovery methods.

**Figure 4.**
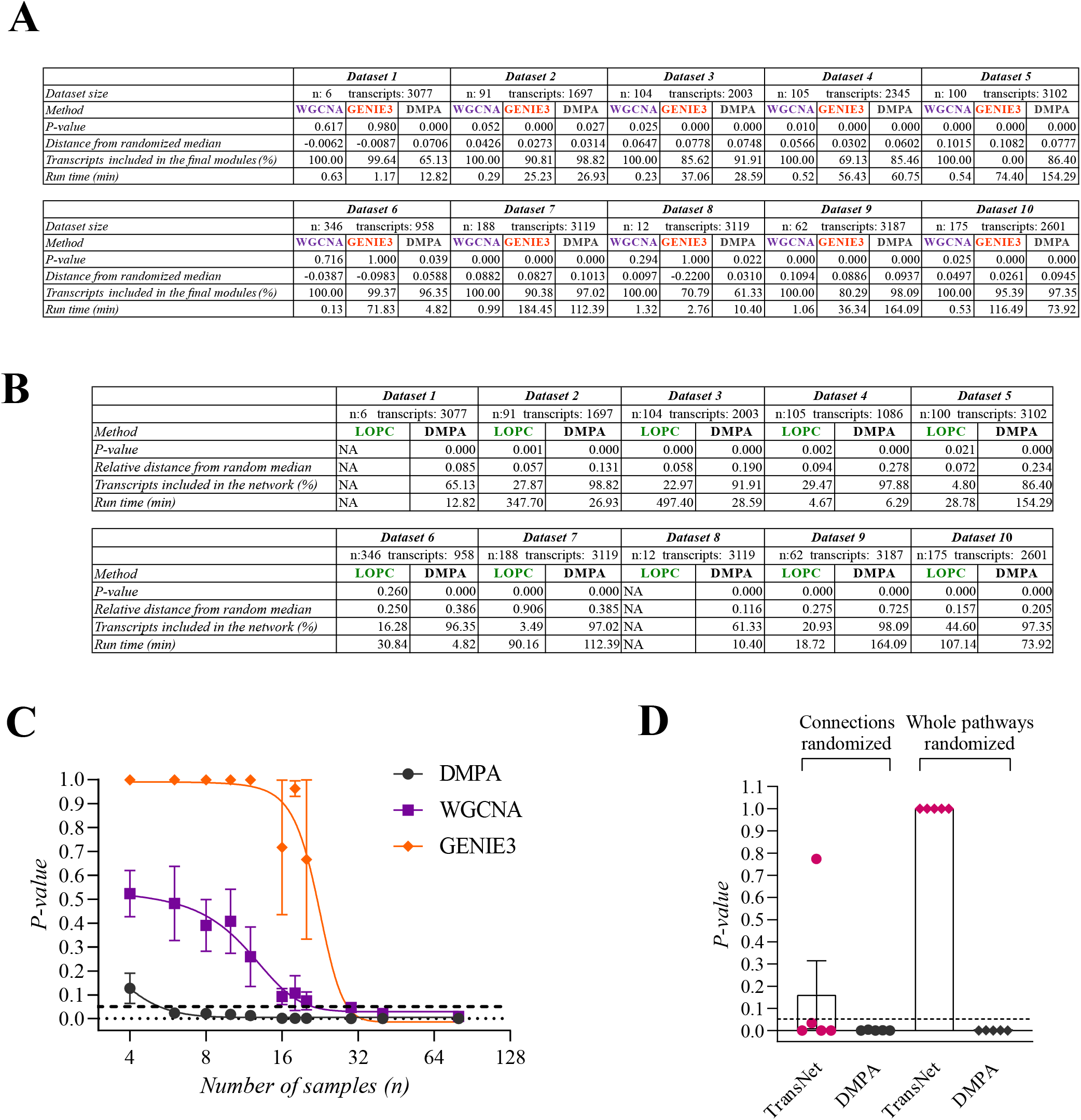
The performance of DMPA against previous methods. **A:** The performance of DMPA against WGCNA and GENIE3 in module discovery. The amount of known transcription factor target gene relationships in transcriptome data, runtime and percentage of transcripts included in the inferred modules was assessed. **B:** The performance of DMPA against LOPC in network discovery. The amount of known transcription factor target gene relationships in transcriptome data, runtime and percentage of transcripts included in the inferred networks was assessed. **C:** The performance of DMPA against WGCNA and GENIE3 in module discovery in transcriptome datasets with different sample sizes. The statistical signficance of the amount of known transcription factor target gene relationships in the inferred modules was assessed. Mean ± SEM. **D:** The performance of DMPA against TransNet in pathway discovery. Multi-omics data was analyzed with both methods and the amount of known associations in pathway annotations from the pathway Commons database in the inferred pathways was assessed. The statistical significance of the amount of the known associations in the inferred pathways was determined with two different statistical methods, where either only the connections between the modules or the whole pathways were randomized. Mean ± SEM.

DMPA was additionally benchmarked against another multi-omics integration method that is independent of prior knowledge. Since DMPA already performed better in module discovery than WGCNA and correlation networks, the examination of the performance of DMPA against these methods with multi-omics integration was considered redundant. Instead DMPA was benchmarked against the closest discovered analogue method, Transkingdom Network Analysis (TransNet) [33]. The aim of TransNet is to discover causal links between two different omics datasets. TransNet first models the omics datasets into correlation networks, assigns dense subnetworks into modules, and then utilizes correlation to find the connecting nodes between the modules of the omics datasets. Available multi-omics datasets were analyzed both with TransNet and DMPA and the ability of the inferred pathways to represent known signaling pathways was estimated as in Figure 3B-C. The ability to connect inferred modules into signaling pathways was also estimated by a version of the validation method utilized in Figure 3B-C where only the connections but not the modules themselves were randomized. As per its aim, TransNet was able to significantly discover connections between the modules that represented known cell signaling pathways, but the modules themselves were not representative of cell signaling pathways as suggested by the lack of a significant P-value from the validation method that also randomized the modules (Figure 4D). DMPA, however, was able to achieve a significant P-value with both versions of the validation method indicating that the pathways inferred with DMPA are closer to known cell signaling pathways.

### Case study: DMPA of cleaved TYRO3 signaling in melanoma cells

To demonstrate the performance of DMPA with a freshly acquired multi-omics dataset, an in-house multi-omics dataset from the cleavage-dependent signaling of the receptor tyrosine kinase TYRO3 was acquired. To this end, a stable TYRO3 knock-down WM-266-4 cell line was created and the cells were transfected with wild-type and cleavage-resistant variants of TYRO3 (ΔGS, ΔADAM) or control vector (Supplementary Figure 12A). The ΔGS variant, resistant to proteolytic cleavage by the gamma-secretase complex, has been described before [34]. The ΔADAM variant, resistant to cleavage by a disintegrin and metalloprotease (ADAM), was created by predicting the ADAM17 of ADAM10 cleavage sites of TYRO3 with a 0th order Markov chain and mutating the key residues in the predicted cleavage sites. The ability of the ΔADAM variant to still act as functional receptor was validated by examining its cleavage (Supplementary Figure 12B), autophosphorylation (Supplementary Figure 12C) and ability to activate a known substrate with western blot analyses (Supplementary Figure 12D). The expected effect of the mutations in the ΔADAM variant on the subcellular location of TYRO3 was examined with immunofluorescence (Supplementary Figure 12E). The analyses strongly suggested that the ΔADAM variant was indeed a functional receptor that was more resistant to cleavage than the wild-type receptor. Interactome, phosphoproteome, proteome and transcriptome data were acquired from the cells expressing the wild-type or cleavage-resistant variants of TYRO3 or the control vector (Supplementary Figure 12F). A differential expression analysis was conducted to discover the differential interactome, phosphoproteome, transcriptome and proteome associated with the full-length of the cleaved intracellular domain (ICD) of TYRO3 (Supplementary Figures 13-18). The details of the methodology to validate the ΔADAM variant and to acquire the multi-omics data is provided in the Supplementary methods.

The differentially expressed signaling events associated with the expression of full-length TYRO3 or the cleaved TYRO3 ICD were analyzed with the DMPA first to extract regulatory complexes. The DMPA discovered 24 previously undescribed interactome, 110 phosphoproteome, 46 transcriptome and 130 proteome regulatory complexes that associated with the expression of full-length TYRO3 (Supplementary Figure 19, Supplementary Table 2). The expression of the cleaved TYRO3 ICD in turn was associated with 53 new interactome, 84 phosphoproteome, 45 transcriptome and 136 proteome regulatory complexes (Supplementary Figure 20, Supplementary Table 2). The inferred regulatory complexes were further combined with the DMPA into pathways, resulting in a total of 51 and 46 unique signaling pathways associating with the signaling stimulated by the full-length and the released ICD of TYRO3, respectively (Supplementary Table 1).

### Enrichment analyses for modules and pathways inferred with the DMPA

To contextualize the function of the modules and pathways inferred with DMPA, an enrichment analysis (please see Materials and methods for details) to predict the upstream transcription factor, upstream kinase, subcellular location and the biological process for each transcriptome, phosphoproteome and interactome module or pathway, respectively, was devised (See supplementary materials and methods for details). The predicted upstream transcription factors, kinases and subcellular locations for the transcriptome, phosphoproteome and interactome modules are provided in Supplementary Tables 3-5. To ensure the validity of the upstream transcription factor and kinase predictions, the predicted transcription factors and kinases were compared to those known to be regulated by TYRO3. Several transcription factors known to be regulated by TYRO3 were predicted to regulate the modeled regulatory transcriptome complexes, such as STAT3 [35], MYC [36,37], and MITF [38] (Supplementary Table 3). Furthermore, the most common transcription factor predicted to regulate the transcriptional complexes associated with TYRO3 signaling was MYC, with the respective gene amplified in the WM-266-4 cell background [39]. Similarly, several kinases reported to be associated with TYRO3, such as AKT [40,41], mTOR [40,42], GSK3 [41,42], MEK1 [42], PKC [42], and CAMK2 [42], were predicted to regulate the phosphoproteome complexes associated with full-length and cleaved ICD of TYRO3 (Supplementary Table 5). The predicted subcellular localizations of the interactome complexes of full-length and the cleaved ICD of TYRO3 (Supplementary Table 4), in turn, were compared to the reported localization difference between the wild-type and the ΔGS and ΔADAM variants of TYRO3 (Supplementary Figure 12E) (Merilahti *et al*, 2017) to ensure the validity of the predicted subcellular locations. The differences in the predicted localizations of the interactome complexes reflected the known difference in the subcellular localization of full-length and cleaved ICD of TYRO3 (Supplementary Table 4). Thirteen out of the 54 (24%) cleaved ICD interactome complexes were predicted to localize into the nucleus, while nuclear localization was predicted for only 2 out of the 21 (8 %) full-length interactome complexes (*P* = 0.032 against frequency of nuclear assignment in the interactome complexes of cleaved TYRO3 ICD). Moreover, 8 out of the 21 (33%) full-length interactome complexes, but only 5 out of the 54 (9%) ICD interactome complexes, were predicted to localize into the plasma membrane (*P* < 0.0001 against frequency of plasma membrane assignment in the interactome complexes of full-length TYRO3). Both the full-length TYRO3 interactome complexes (5 out of 21; 20%) and the ICD interactome complexes (11 out of 54, 20%) were equally predicted to be localized in the cytoskeletal structures (*P* = 0.0025 for full-length TYRO3 and *P* < 0.0001 for TYRO3 ICD against frequency of cytoskeleton assignment in randomly modelled interactome complexes).

The summary of the predicted functions for each pathway associated with the signaling of full-length and ICD of TYRO3 is displayed in Figure 5 and Supplementary Figure 21. The functional enrichment analysis of the modelled pathways of full-length and ICD of TYRO3 was able to identify unique pathways associated with various cellular processes including ones involved in the pathogenesis of cancer. A significant difference was noted in the number and direction of pathways related to growth and cell cycle, cell adhesion and motility, cell morphology, cell death, and immune response between the cells expressing the full-length versus the ICD of TYRO3 (Figure 5). These findings indicate that a differential response to these functions could be enacted by the full-length and ICD of TYRO3.

**Figure 5.**
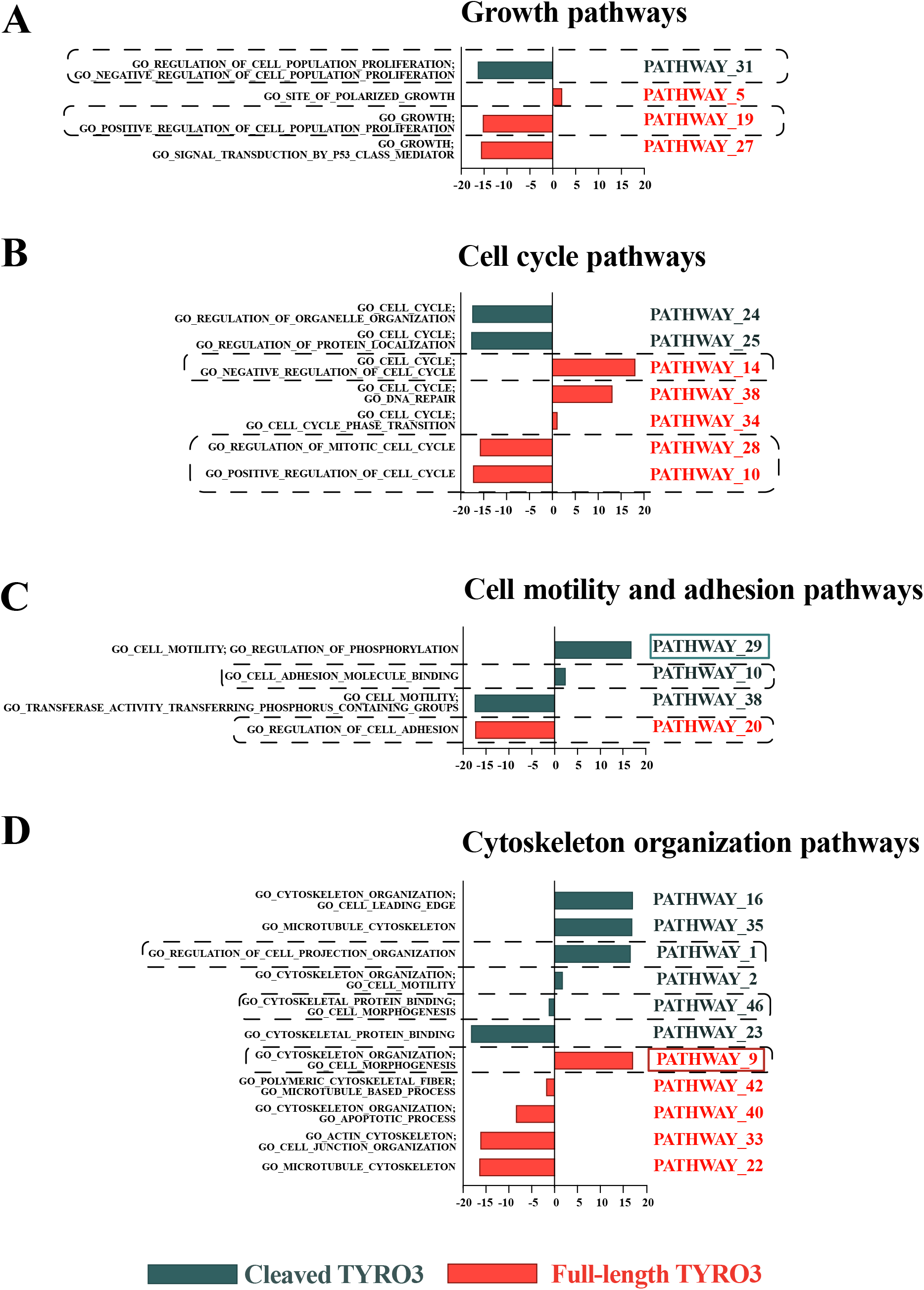
The functional categorization of the selected full-length and cleaved TYRO3 pathways inferred with the DMPA. The predicted function of the multi-omics pathways of the full-length and the cleaved ICD of TYRO3 inferred with the DMPA. The bars represent the median pseudolog2 fold change of all the proteins, transcripts and phosphorylated residues in the pathway against the control condition. For a full list of predicted functions for the pathways of full-length and ICD of TYRO3 see Supplementary Figure 21. For further details on the pathways see supplemental tables 1 and 2. The cell behavior predicted by the pathways surrounded with the dashed line is examined in Figure 6A-C.The highlighted pathways are visualized in Figure 6D-E.

### Functional validation of the functional enrichment analyses of the DMPA

To validate the ability of function enrichment of the DMPA to predict differential cellular functions based on molecular data, we randomly chose 3 cellular functions predicted to go to opposite directions by the signaling pathways activated with the ICD or the full-length TYRO3. First, to assess proliferation induced by the ICD or the full-length TYRO3, the growth of WM-266-4 transfectants was analyzed using live-cell imaging (Figure 6A). The function prediction of the ICD pathway 31 and the full-length pathways 19 and 10 (Figure 5A-B) indicated that the cleavage and release of the ICD of TYRO3 would promote cellular growth, while the expression of the full-length TYRO3 would down-regulate the positive regulation of proliferation. Indeed, the WM-266-4 cells expressing wild-type TYRO3 proliferated significantly faster than the vector control cells or cells expressing either the ΔGS or the ΔADAM variants of TYRO3 (Figure 6A).

**Figure 6.**
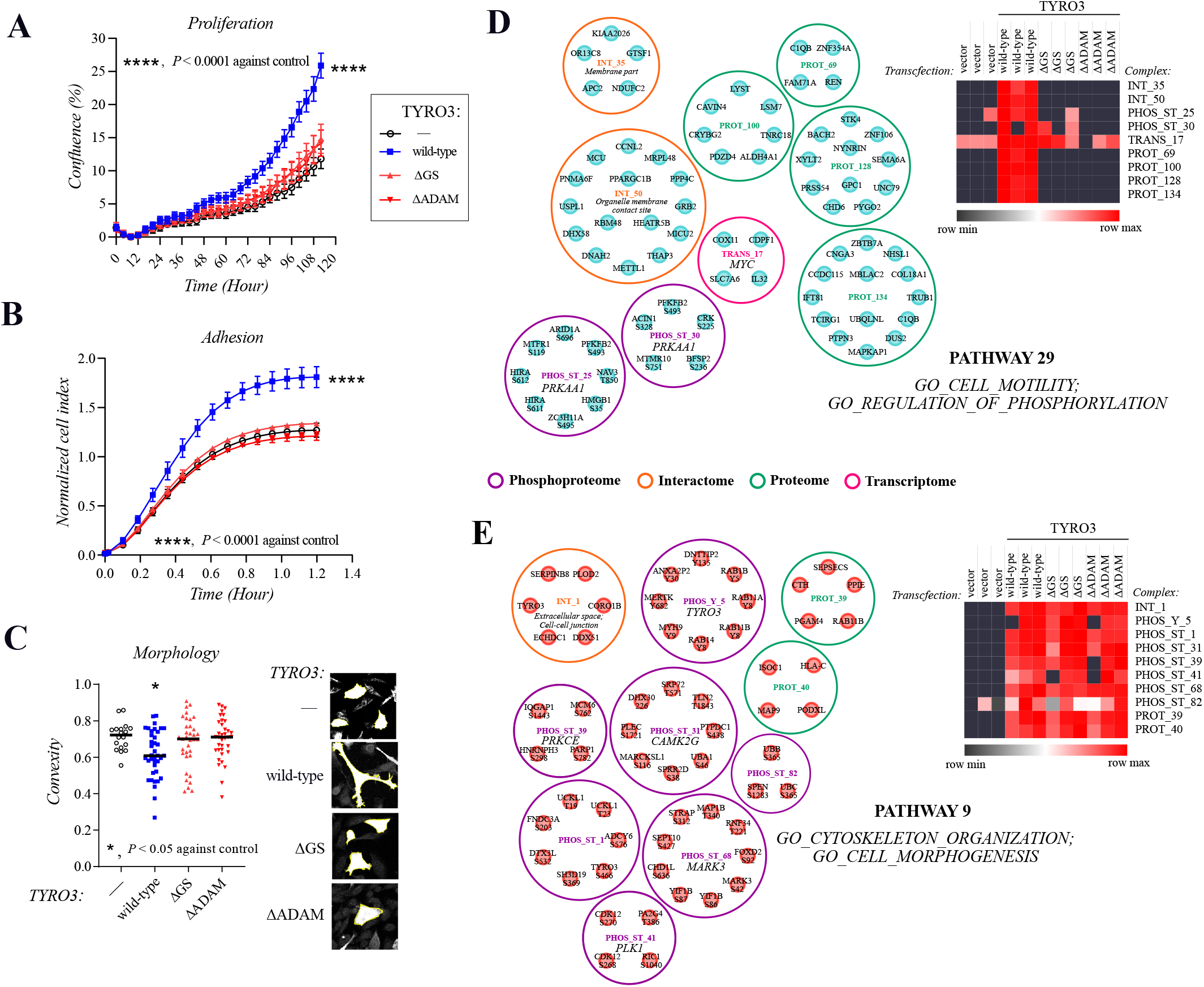
Functional validation and composition of the selected pathways of full-length and cleaved ICD of TYRO3. **A:** Proliferation of WM-266-4 transfectants was measured with live-cell imaging. For statistical testing, the non-parametric Friedman 2-way ANOVA and the Dunn’s multiple comparison test with a multiple test correction for the P-values was utilized. The symbols represent the mean and the whiskers the standard deviation of the values. A representative plot from 1 out of 3 independent replicate experiments is shown (n=6). **B:** Real-time adhesion of WM-266-4 transfectants was explored with the xCELLigence cell impedance measurement system. For statistical testing, the parametric 2-way ANOVA and the Dunn’s multiple comparison test with a multiple test correction for the P-values was utilized.The symbols represent the mean and the whiskers the standard deviation of the values. A representative plot from 1 out of 4 independent replicate experiments is shown (n=5). **C**: The morphology of WM-266-4 transfectants was analyzed from thresholded confocal images taken in plane with the plasma membrane with MorphoLibJ plugin of ImageJ. For statistical testing, the non-parametric Kruskal-Wallis ANOVA was utilized. The post hoc analyses were conducted with the Mann-Whitney U test and the resulting P-values were corrected with the method of Benjamini, Krieger and Yekutieli. Convexity: the ratio between the convex perimeter and the real perimeter. One dot represents one cell and the horizontal line the median value. Combined results from 4 indenpendent experiments is shown. **D-E:**Visualization of the cell motility pathway 29 of cleaved TYRO3 ICD (**D**) and the cytoskeleton organization pathway 9 of full-length TYRO3 (**E**). The median expression of all the transcripts, proteins or phosphorylated residues in the indicated samples and regulatory complexes is presented in the heatmaps. Individual complexes are separated by circles. The inferred regulatory complexes are identified by a letter combination (INT for interactome, PHOS_ST for serine/threonine phosphoproteome, PHOS_Y for tyrosine phosphoproteome, TRANS for transcriptome, PROT for proteome) and a number. Predicted transcription factors, kinases and subcellular locations are indicated with italic letters under the complex identifier.

As another read-out to validate the functional enrichment analyses of the pathways inferred with DMPA, the adhesion rate of the WM-266-4 transfectants to fibronectin coated wells was investigated by real-time cellular impedance measurement (Figure 6B). Again, as predicted by the ICD pathway 10 and the full-length pathway 20 (Figure 5C), the cells expressing wild-type TYRO3 adhered to fibronectin with a greater affinity than vector control cells or cells expressing either the ΔGS or ΔADAM variant of TYRO3 (Figure 6B).

Finally, the two-dimensional cellular morphology of the WM-266-4 transfectants were examined from thresholded confocal images with the ImageJ MorphoLibJ plugin (Figure 6C). As indicated by the ICD pathway 46 and the full-length pathway 9 (Figure 5D), the cells expressing wild-type TYRO3 exhibited distinct morphology from the vector control cells or cells expressing either the ΔGS or ΔADAM variant of TYRO3 (Figure 6C). The morphological difference was captured by the convexity measure (the ratio between the cell area and its convex area), the lower value of which suggests a shape with more protrusions. A greater amount of cell protrusions in the cells expressing wild-type TYRO3 was also predicted by the DMPA (ICD pathway 1 in Figure 5D). Furthermore, the WM-266-4 cells expressing wild-type TYRO3 or either of the TYRO3 mutants demonstrated less circular cell shape than the vector control cells, indicating that the full-length TYRO3 additionally regulates cell morphology as suggested by the full-length pathway 9 (Supplementary Figure 22). Taken together, these validation experiments suggest that the function enrichment of the pathways modeled with the DMPA can predict both the function and the direction of the function from purely molecular multi-omics data.

## Discussion

The increased demand for a ground-up analysis method able to model multi-omics data into cell signaling pathways motivated us to create the de novo multi-omics pathway analysis, DMPA. The basis of the DMPA lies in the calculation of two robust metrics of molecular association, the correlation and stoichiometry score, and the consequent modeling of the cell signaling molecules into regulatory complexes based on the nearest neighbor principle. The DMPA was able to consistently find relevant biological associations in both published and freshly acquired multi-omics data. The method showed consistent performance and the combined score-based networks outperformed correlation-based network models in all tested data types. DMPA outperformed previous methods in both module and pathway discovery.

The advantages of the DMPA lie in its independence on previous knowledge, robustness, which makes it easily scalable to other omics data types, low computational demand and ability to work with low sample sizes. The proof of the predictive ability to discover relevant molecular association opens up further applications for the combined score in more sophisticated and complex inference methods. Previous multi-omics integration methods have been focused mostly on feature extraction and matrix factorization strategies. While the inferred latent factors have been found accurate in stratifying samples into clusters and can be predictive of clinical outcome, there is little evidence that the discovered factors would represent actual signaling pathways in the cell [10,11]. Here we provide evidence that the pathways modelled with the DMPA reflect previously described signaling pathways.

Since the DMPA is based on the inherent variation between samples in the omics data, the quality of the omics data poses a limitation to the analysis. Data wrought with technical variation, or inflated with missing values or lack of repetitions, will lead to loss of model accuracy. The nonparametric formulation and a zero inflated version of the stoichiometry score was devised to battle two of these limitations. Since the correlation, stoichiometry and combined scores are ranked and thus dependent on the choice of the signaling molecules included in the dataset, a choice of the cut-offs for the data to be modeled will also significantly affect the final modeled pathways. Involving or omitting proteins, post-translational modifications or transcripts from the omics data used for the analysis will influence the inferred final pathways. For accurate inference of the multi-omics pathways, the multi-omics data should be acquired from the same samples. The DMPA does not support combination of multi-omics data that have been acquired from different samples even though they would represent the same tissue type or cell line.

We acquired multi-omics data from the cleavage-resistant and wild-type TYRO3 variants and utilized the DMPA to shed light on the unknown cleavage-associated signaling pathways of the TYRO3 RTK in melanoma cells. In addition to signaling as a full-length RTK, TYRO3 has been proposed to undergo regulated intramembranous proteolysis (RIP) by sequential proteolysis by the gamma-secretase complex followed by cleavage by an ADAM protease, releasing a soluble ICD with potential signaling activities [34]. The DMPA discovered a total of 51 and 46 differential unique signaling pathways for the full-length TYRO3 and the cleaved TYRO3 ICD, respectively. An enrichment analysis to infer the biological process the pathways were involved with was also devised. A difference in the amount and the direction of the pathways associated with growth and cell cycle, cell motility and adhesion, immune response, chromosome organization, cell death, and cell differentiation were observed between the full-length and cleaved TYRO3 ICD. Many of these functions may be critically involved e.g., in the progression or therapeutic responses in melanoma. For example, TYRO3 has been implicated in the proliferation, tumorigenesis, chemoresistance and motility of melanoma cells [38,43–45]. Validation experiments confirmed that the DMPA with the function prediction was able to accurately predict both the function and the direction of the function of the cells expressing either wild-type or cleavage-impaired variants of TYRO3. These findings also represent the first comprehensive characterization of signaling pathways stimulated by the gamma-secretase-released ICD of a RTK as compared to the canonical signaling *via* the full-length receptor.

The DMPA segregates all signaling events into pathways independent on the amount of previous knowledge on the modeled molecules. This feature is crucial in discovery of novel signaling modules. Since the DMPA can predict molecular associations, it can predict upstream transcription factors of transcripts, kinases of phosphorylation sites, and subcellular locations of proteins in the regulatory complexes, for which these features are unknown. Similarly, the DMPA can predict connections to biological functions for proteins, transcripts or post-translationally modified proteins in the pathways for which no previous connection to the biological function has been discovered. The pathways inferred with DMPA can be re-entered into DMPA to acquire super pathways and thus broader connections between the modelled pathways can additionally be discovered with the DMPA.

## Materials and methods

A more comprehensive description of the materials and methods can be found in the Supplementary Materials and Methods.

### The inference of regulatory complexes and their combination in the DMPA

Matlab R2016a (MathWorks) was used as the coding environment. The pair-wise association between two proteins, phosphosites or transcripts was determined by using a combined score. The combined score is derived from the multiplication of the correlation and the stoichiometry score the mathematical formulations of which are presented in Figure 1A. The correlation score for all the possible protein, phosphosite or transcript combinations was calculated with the Spearman correlation, ranked and adjusted to derive scores from 1 to 0 in equal increments. The stoichiometry score for all the possible protein, phosphosite or transcript combinations was calculated by dividing the third quartile value (Q3) of relative abundances with the first quartile (Q1) value and similarly ranked and adjusted to derive scores from 1 to 0 in equal increments. Two versions of the stoichiometry score were devised for non-zero-inflated and zero-inflated data. In the non-zero-inflated version of the stoichiometry score, relative abundance data were considered only from samples in which the expression value for both intracellular molecules or their states was non-zero. In the zero-inflated version of the stoichiometry score (weighted stoichiometry score), the stoichiometry score is punished (inflated) if one of the two intracellular molecules or their states in the same sample has a missing value. The stoichiometry score, in turn, is rewarded (deflated) if both intracellular molecules or their states have missing values in the same sample.

Cut-off parameters C and S were devised to reduce computational time to only consider combined scores higher than a certain threshold. The cut-off parameter C is determined by the minimum number of interactions considered for intracellular molecules that can be set in the analysis. The cut-off parameter S is determined by the number of intracellular molecules or their states in the dataset that are estimated to not have any association.

The regulatory complex inference analysis was designed to utilize the nearest neighbor concept to find the highest scoring (sum of the combined score between all 3 members) 3 member networks for all intracellular molecules in the list and to combine and trim the networks based on common network members and combined scores as described in the following order: 1) Networks with 2 common members are combined until no two complexes with 2 common members can be found. 2) Networks with only one common member are combined if the size of both networks is under or equal to the set parameter value and the network score is higher or equal to 2 indicating a strong association. The parameter for this joining (parameter 6 in regulatory complex inference, maximum size of a complex to be joined with only one common node) can be set if smaller regulatory complexes are preferred. 3) Networks with 2 common members are again combined until no two complexes with 2 common members can be found. 4) Two out of the 3 members of the remaining 3 member networks are removed if 2 out of three members are already present in another complex. 5) The remaining 3 member networks are joined if they still have a common member. This step can be omitted if smaller or larger regulatory complexes are preferred (parameter 7 in regulatory complex inference). The analysis allows for an intracellular molecule or their state to be a member for more than one regulatory complex.

The same strategy was used for the combination of the regulatory complexes into larger modules as was the derivation of the regulatory complexes in the first place with a few modifications. The median value of all complex members in each sample was used to combine the regulatory complexes and the absolute value of the Spearman correlation was used to allow equal combination of up-and downregulated regulatory complexes. The default settings were used for the analysis of the TYRO3 signaling pathways.

### Validation of regulatory complexes and their combination

The data on known protein-protein interactions, kinase substrate phosphosite relationships, transcription factor target gene relationships, methylation trait relationships and metabolic reactions were acquired from the STRING, PhosphoSitePlus, ChEA3, and Rhea databases, respectively [26– 29,46,47]. The data on annotated signaling pathways was acquired from PathwayCommons [31]. The validation interactome, phosphoproteome, transcription and metabolomics data were acquired from the publications of Bath et al and Karayel et al, and from ArchS4 and MetaboLights database, respectively [24,25,48,49]. The methylation data and multi-omics data for the regulatory complex combination to pathways was acquired from the LinkedOmics database [30], where a combination of either proteomics, transcriptomics, methylomics, phosphoproteomics, protein array or acetylation data from the same samples were analyzed and combined. The sum of all protein-protein interactions, kinase substrate phosphosite relationships or transcription factor target gene relationships in the modeled regulatory complexes was used as the validation score for interactome, phosphoproteome and transcriptome regulatory complex validation, respectively. The sum of co-occurrences of two complex members from two different combined complexes in the same pathway annotation for all the combined complexes was used as a validation score for regulatory complex combination. To validate the regulatory complexes, the validation score from the modelled regulatory complexes was compared to the sum of all protein-protein interactions, kinase substrate phosphosite relationships or transcription factor target gene relationships in randomly modeled regulatory complexes of same size to derive an empirical probability density function from 1000 rounds of simulation. To validate the combination of regulatory complexes into pathways, the validation score from the modelled pathways were compared to the sum of all co-occurrences of two complex members from two different randomly modelled and combined complexes of same size in the same pathway to derive an empirical probability density function from 1000 rounds of simulation. An Epanechnikov kernel was used to fit the simulated data into the empirical probability density function. The corresponding P-values were drawn from the empirical cumulative distribution function. The relative distance of the validation score from the median of the empirical probability distribution was calculated by subtracting the median value from the validation score and by dividing the remainder with the median value. To assess the effect of different scores on the accuracy of the modeled regulatory complexes and pathways, the complexes were modeled with either correlation score, stoichiometry score or the combined score and combined with either correlation score, stoichiometry score or the combined score. The conservation of the different scores was assessed in each data type by deriving the median score of the modeled complexes in different datasets from the one used for the initial modeling.

### Benchmarking

The same datasets that were used for the validation of the regulatory complexes and pathways inferred with DMPA were used for benchmarking. The R implementations of WGCNA [3], LOPC [50], GENIE3 [32] and TransNet [33] was used according to developers instructions. The power for the WGCNA analysis was set based on the scale free topology model fit and mean connectivity as instructed by the WGCNA documentation. The genes predicted to be controlled by the same upstream gene in GENIE3 were considered as part of the same module. The validation score and statistical significance of the validation score was estimated as for the regulatory complexes for modules derived with WGCNA and GENIE3. Since LOPC only returns network edges, a separate validation strategy to compare the performance of LOPC and DMPA was developed. The amount of known transcription factor target gene relationships in the network edges returned by LOPC as determined by the ENCODE database [28] was calculated and used as a validation score. The validation score was compared to the sum of all transcription factor target gene relationships in randomly modeled network of same size to derive an empirical probability density function from 1000 rounds of simulation. An Epanechnikov kernel was used to fit the simulated data into the empirical probability density function. The corresponding P-values and relative distance from the random median were estimated from the empirical cumulative distribution function. To compare DMPA against LOPC the amount of known transcription factor target gene relationships in the final edges in the regulatory complexes as suggested by the remaining combined scores after cut-off and trimming were considered as network edges. The validation score and the empirical probability distribution were similarly estimated for the networks suggested by DMPA. The validation score for the pathways suggested by the TransNet analysis were estimated as those for the DMPA. The clusters connected by the connected nodes were considered a pathway. A version of the pathway validation method where only the connections between different modules were randomized was developed and applied to both TransNet and DMPA.

### Immunofluorescence and confocal microscopy

To detect ectopically expressed V5-tagged TYRO3 constructs, WM-266-4 transfectants were cultured on coverslips in serum-free conditions overnight. The cells were fixed with 3% paraformaldehyde, and permeabilized with 0.1% Triton X-100. Cells were stained with anti-V5 (13202; Cell Signaling Technologies) and AlexaFluor 555 goat anti-rabbit (Molecular Probes). After labeling the nuclei with 4’,6-diamid-ino-2-phenylindole (DAPI; Sigma-Aldrich), the cells were mounted with Mowiol 40-88 (Sigma-Aldrich). Images were acquired with Zeiss LSM 880 confocal microscope (Zeiss). Morphological analysis were carried out from confocal slices in plane with the plasma membrane using MorphoLibJ v 1.4.2.1 [51] plugin in Image J.

### Live-cell imaging of cell growth

The WM-266-4 cell transfectants were plated in serum-free medium at 30,000 cells/24-well. The wells were imaged every 2 hours for 120 hours and cell confluence was measured with Incucyte ZOOM (Sartorius).

### Real-time cell adhesion assay

The WM-266-4 cell transfectants were detached using 5 mM EDTA 6 to 12 hours after transfection. The cells were plated at 15,000 cells/well onto fibronectin-coated (5 μg/cm^2^) xCELLigence E-plate (Agilent) wells in serum free medium. Real time cell impedance was measured with the xCELLigence RTCA analyzer (Agilent) for 24 hours. The resulting cell index values were normalized to the number of cells remaining in the wells after 24 hours.

### Information visualization and statistics

For statistical analysis, the GraphPad Prism software (Version 9.02; GraphPad Software) and Matlab 2016a (MathWorks) were utilized. The details of the statistical testing are described in figure legends. All datasets were tested for normality and parametric or non-parametric testing was used accordingly. Heatmaps were generated with Morpheus (https://software.broadinstitute.org/morpheus). The visualizations of the inferred regulatory complexes and pathways were created with Cytoscape (Version 3.8.2; Shannon *et al*, 2003) and Affinity Designer (Version 19.0; Serif)

### Data availability

The mass spectrometry proteomics data have been deposited to the ProteomeXchange Consortium via the PRIDE partner repository with the dataset identifier PXD028881. Reviewer account details: Username: Password: 9e92pZvD. The RNAseq data have been deposited to the Gene Expression Omnibus database with the identifier: GSE190431. The reviewer token (password) to access the dataset in GEO: qxohymmyfvivdyp. The DMPA is available both as matlab code and executables in Mendeley Data: https://data.mendeley.com/datasets/m3zggn6xx9/draft?a=71c29dac-714e-497e-8109-5c324ac43ac3.

### Source data files

The raw source data for the western blots in Supplementaty Figure 12 are available in Supplemenrary Figure 12 – source data 1.zip file. The zipped folder includes both the raw images and the Supplemenrary Figure 12 Annotated Immunoblots.pdf file where the bands presented in Supplemenrary Figure 12 are annotated and indicated in the full gel images. The raw source data for Figure 6A and B is available in the Figure6_source_data.xlsx file that includes the numerical values of all replicate experiments of Figure 6A and B.

## Acknowledgements

We thank Maria Tuominen for skillful technical assistance. This work was supported by the Academy of Finland, Cancer Foundation Finland, the Sigrid Juselius Foundation, the Turku University Central Hospital, Orion Research Foundation, K. Albin Johanssons Foundation, the Cancer Society of Southwestern Finland, the Jenny and Antti Wihuri Foundation, the Maud Kuistila memorial Foundation, Emil Aaltonen Foundation, the Finnish Foundation for Cardiovascular Research, the Paavo Nurmi Foundation, the Aarne Koskelo Foundation, the Paulo Foundation, Finnish Cultural Foundation and the Varsinais-Suomi Regional Fund of the Finnish Cultural Foundation.

## Author contributions

JAMM, KV, KE designed the study, JAMM, KV, and VKO performed the experimentation. KV created the DMPA. TYRO3 knock-down cells were prepared by VKO. KV and KE wrote the manuscript with the help of JAMM.

## Abbreviations used

ADAM: a disintegrin and metalloprotease;
ICD: intracellular domain;
RTK: receptor tyrosine kinase;
DMPA: de novo multi-omics pathway analysis;
WGCNA: weighted correlation network analysis;
LOPC: low-order partial correlation,
TransNet: Transkingdom Network Analysis;

