## Supplementary Figures 1 to 22 for "De Novo multi-omics pathway analysis (DMPA) designed for prior data independent inference of cell signaling pathways"

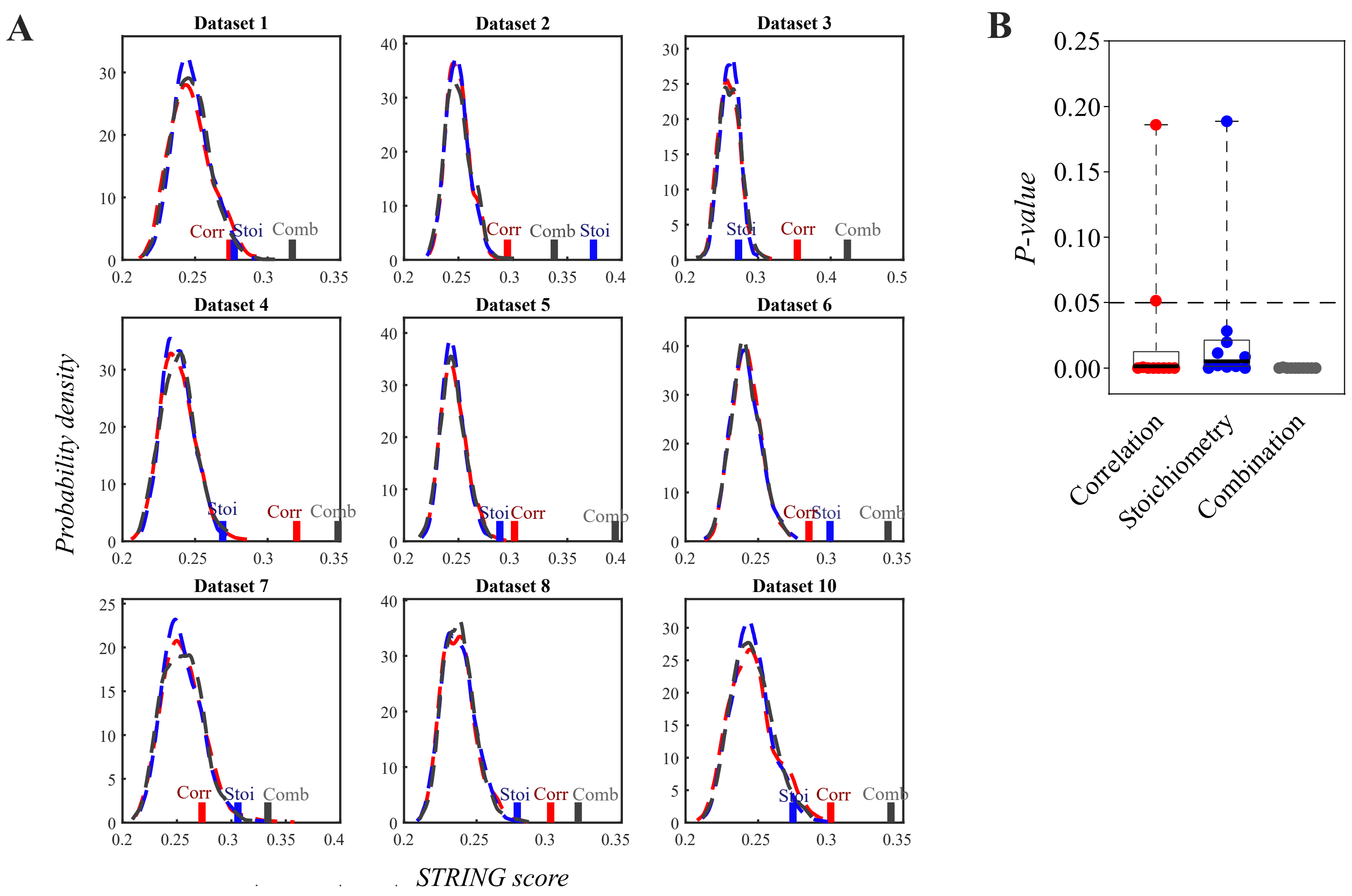


**Supplementary Figure 1. Additional validation of regulatory interactome complexes modeled with DMPA.** Empirical probability densities (**A**) simulated by randomizing the proteins of the modeled regulatory interactome complexes into complexes of the same size and the corresponding P-values (**B**) for the complexes modeled from interactome data by either correlation (Corr, red), stoichiometry (Stoi, blue) or combined score (Comb, grey). The median STRING score of the association between two proteins inside the modeled complexes as determined by the STRING protein-protein interaction database were used as a validation score. The empirical probability densities are visualized with dashed lines of the corresponding color and the score for the modeled network in solid line. One dot in the boxplot represents a P-value for one dataset and the line the median value. The datasets were acquired from published interactome data.

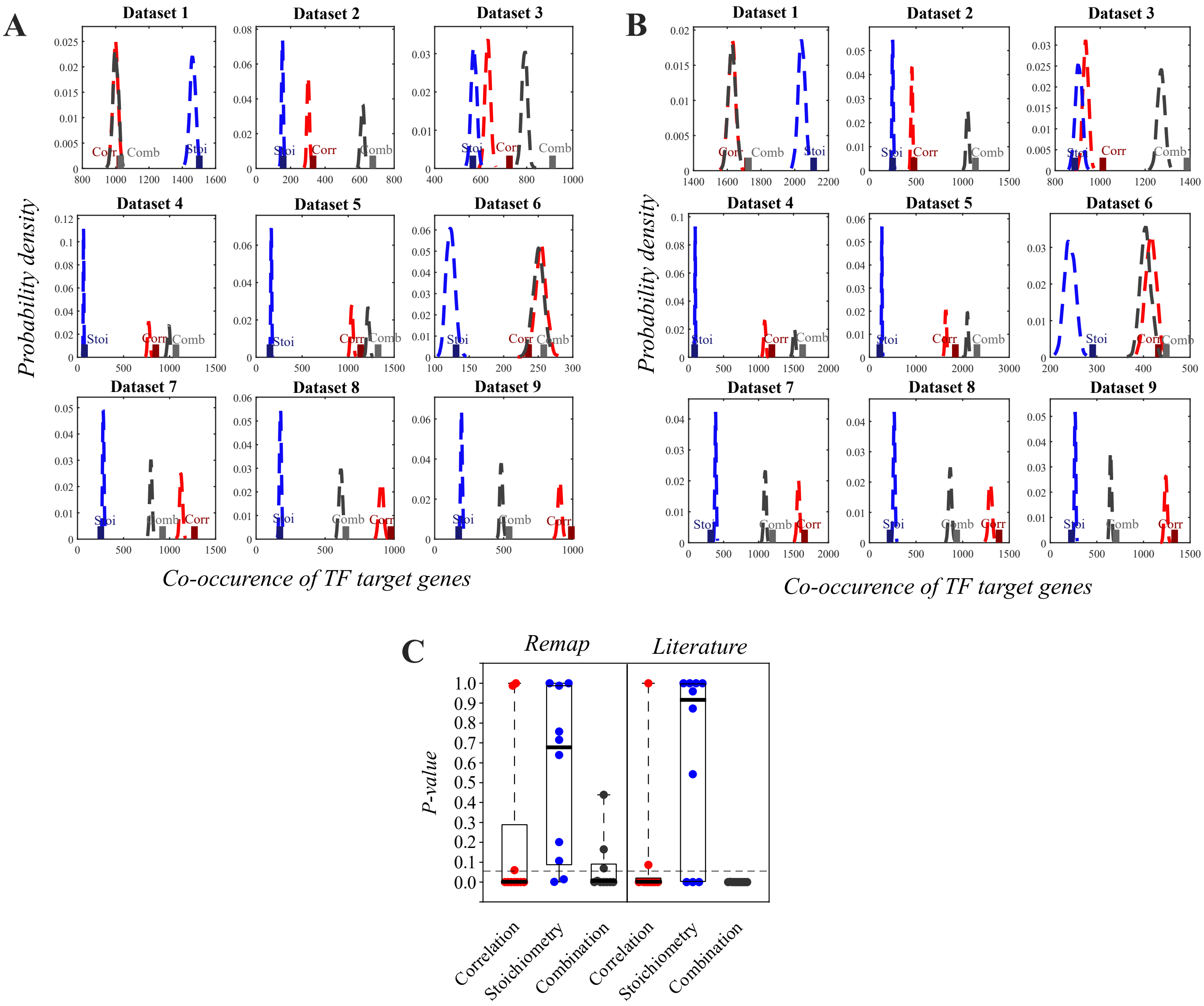


**Supplementary Figure 2. Additional validation of regulatory transcriptome complexes modeled with DMPA.** Empirical probability densities (**A, B**) simulated by randomizing the transcripts of the modeled regulatory transcriptome complexes into complexes of the same size and the corresponding P-values (**C**) for the complexes modeled from transcriptome data by either correlation (Corr, red), stoichiometry (Stoi, blue) or combined score (Comb, grey). The sum of co-occurrences of transcription factor target genes as determined by the Remap (**A**) and Literature (**B**) databases in the modeled complexes were used as a validation score. The empirical probability densities are visualized with dashed lines of the corresponding color and the score for the modeled network in solid line. One dot in the boxplot represents a P-value for one dataset and the horizontal line the median value. The datasets were acquired from published transcriptome data.

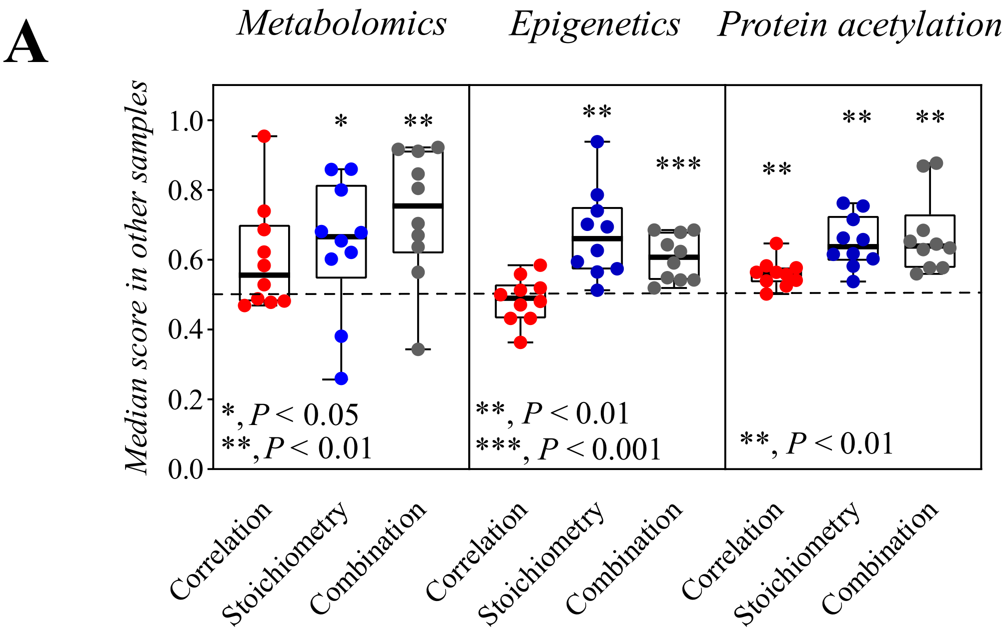


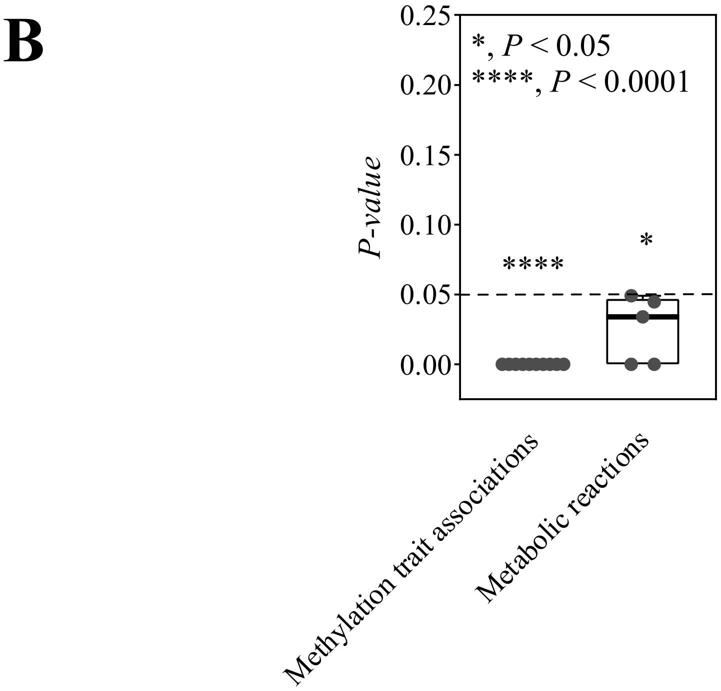


**Supplementary Figure 3. Validation of regulatory complexes inferred with DMPA with additional omics data.**
**A:** Conservation of the indicated scores in metabolomics, methylation and protein acetylation data. Two-tailed one-sample T-test.
**B:** Ability of the regulatory complexes inferred with DMPA to predict methylation trait associations and metabolic reactions in epigenetics and metabolomics data, respectively. The combined score was used. One-tailed one sample Wilcoxon rank test.
The metabolomics data was acquired from the Metabololights database. The methlyation and protein acetylation data was acquired from the CPTAC data accessed through the LinkedOmics database.

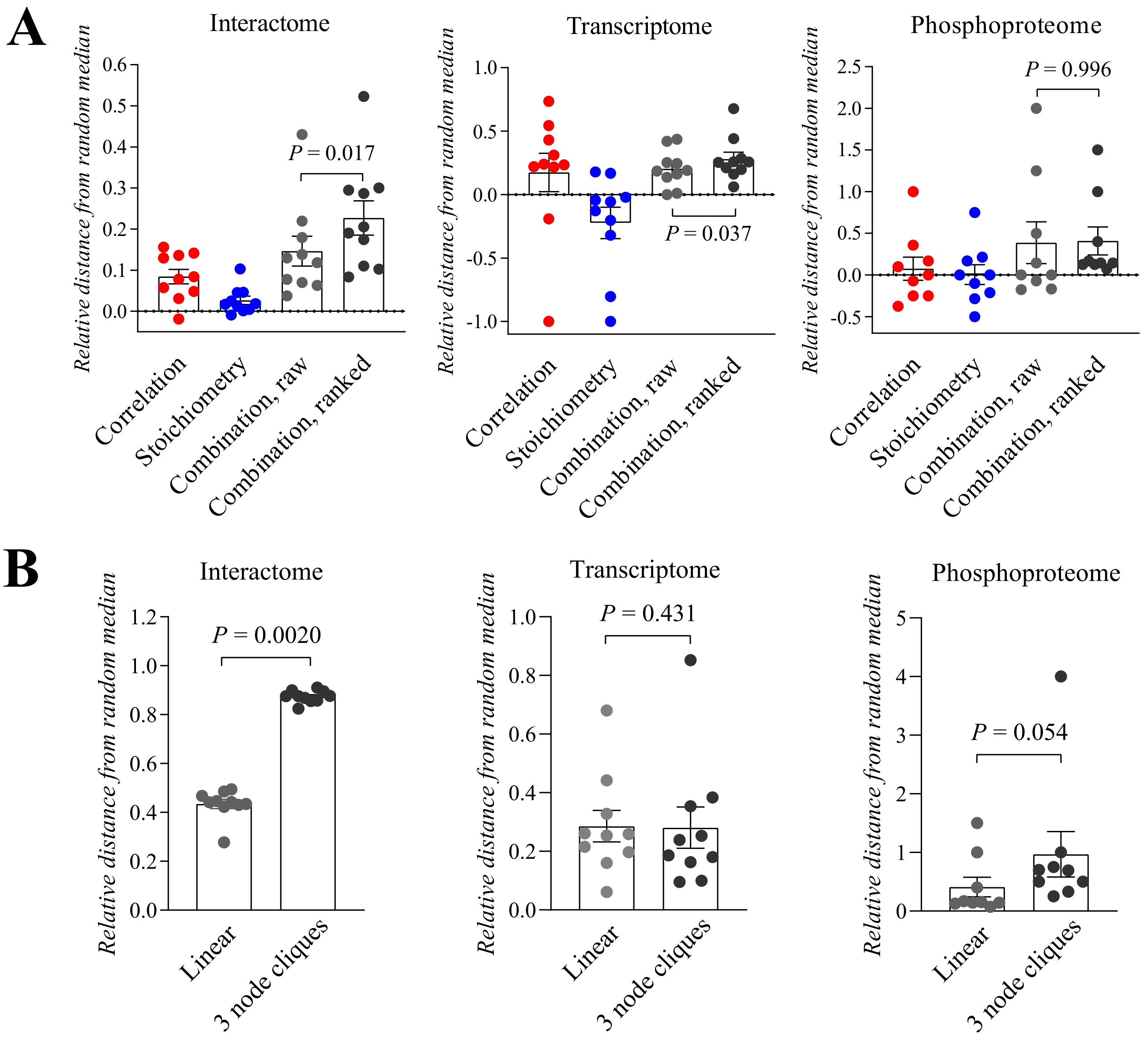


**Supplementary Figure 4. Validation of the design decisions of DMPA.**

**A:** The effect of ranking and adjusting the stoichiometry score and correlation score on the accuracy of DMPA to discover known molecular associations in interactome, transcriptome and phosphoproteome data. The correlation, stoichiometry, raw combination and ranked combination score were calculated and the maximal scoring pairs for each intracellular molecule in each condition was determined. The sum of co-occurences of the maximal scoring pairs in protein-protein interaction , transcription factor target gene and kinase substrate annotations as determined by the STRING, ENCODE and PhosphoSitePlus databases were calculated. Randomized empiricial probability densities were estimated by calculating the sum of co-occurences for random pairs. The distance of the sum of the co-occurences from the median of the random density was estimated in each condition. The distance was normalized against the median of the random density. Repeated measures one-way ANOVA with Dunnett's multiple comparisons test.

**B:** The effect of assigning intracellular molecules or their states to maximally scoring 3 node cliques instead of a linear approach on the accuracy of DMPA to discover known molecular associations in interactome, transcriptome and phosphoproteome data. In the linear approach one maximally scoring interactor was assigned to each intracellular molecule or their states. Relative distance of the validation score from the randomized median is shown.Wilcoxon matched-pairs signed rank test.

**A**

| | Parameter 7 | Parameter 8 |
|---|---|---|
| **Setting 1** | 4 | 5 |
| **Setting 2** | 4 | 0 |
| **Setting 3** | 0 | 5 |
| **Setting 4** | 0 | 0 |

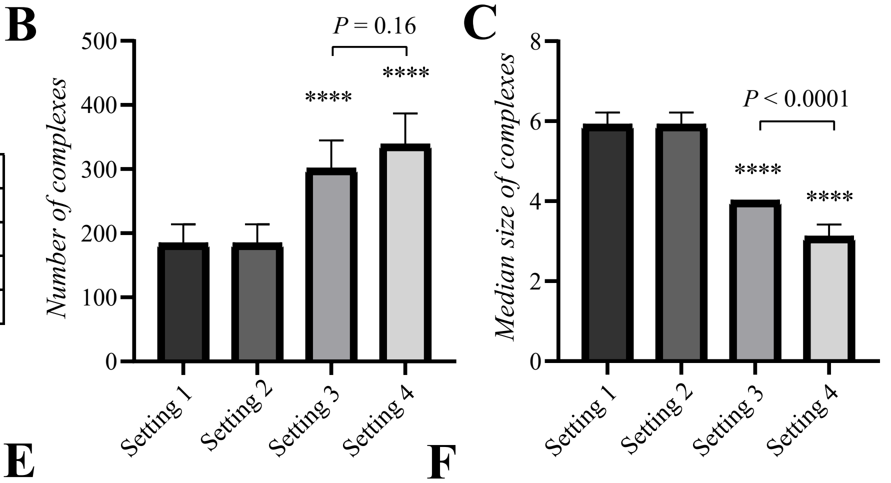


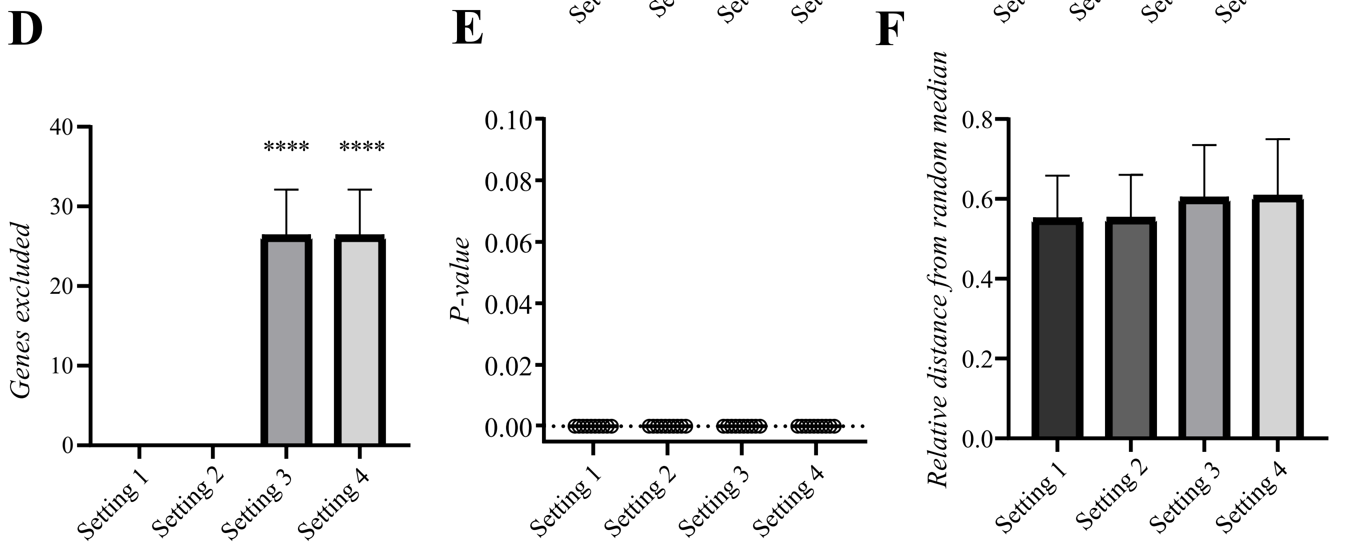


**Supplementary Figure 5. Effect of the size parameters on the inferred regulatory complexes and performance of DMPA.**
**A:** Parameter values for the size parameters 7 (Maximum size of a complex to be joined with only one node) and 8 (Maximum combined size of two small complexes to be joined in the last round) in regulatory complex inference in different settings.
**B:** Number of complexes in the models inferred with different settings.
**C:** Size of regulatory complexes in the models inferred with different settings.
**D:** Number of genes excluded from the models inferred with different settings.
**E-F**: P-value and the distance from the median of the randomized probability distribution of the validation score of the models inferred with different settings. ****, $P < 0.0001$ against Setting 1. One-way ANOVA and Tukey's multicomparison test was used for statistics. Mean ± SD (n=10).

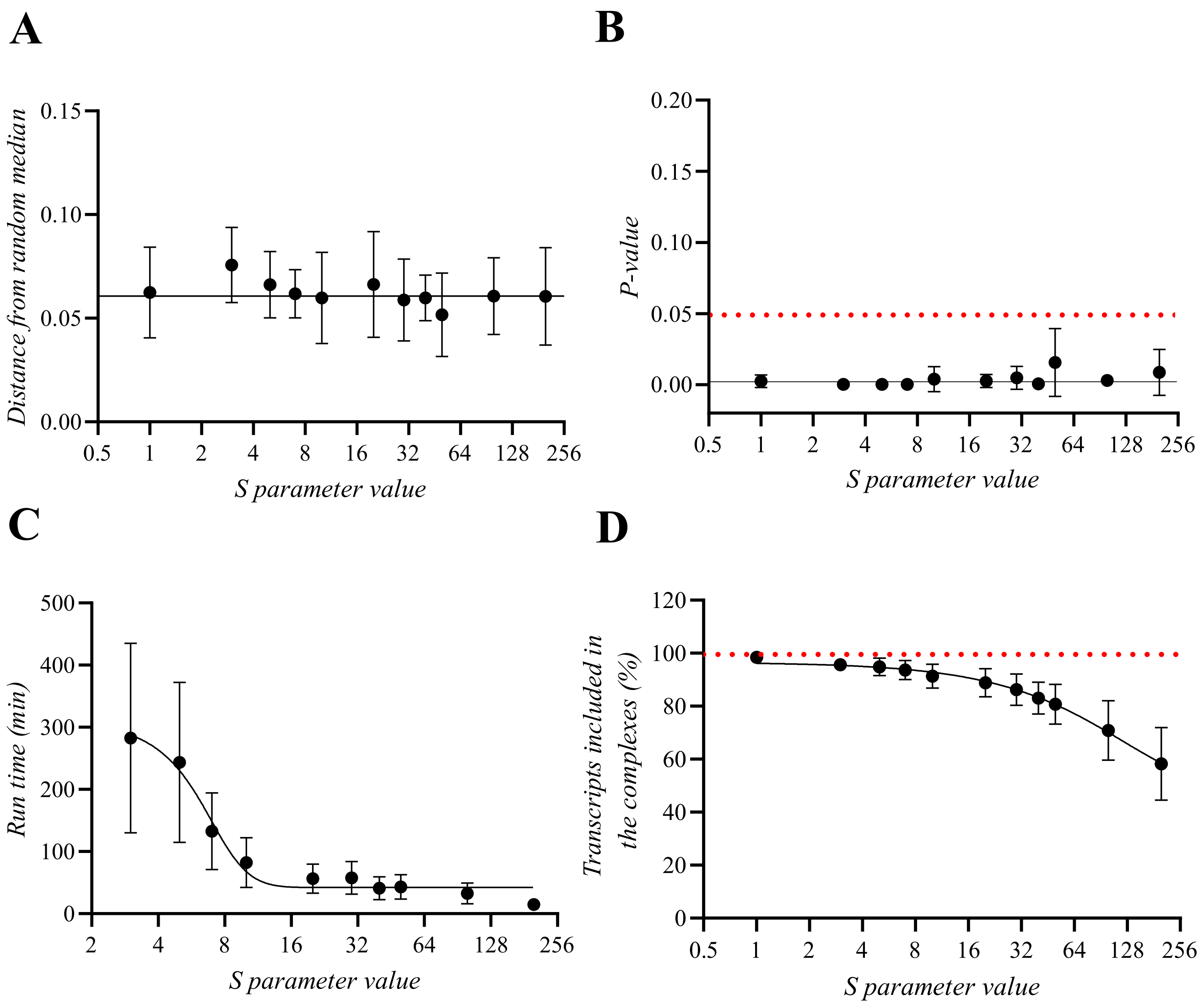


**Supplementary Figure 6. Effect of parameter S choice on the performance of DMPA with transcriptome data.**
**A-B:** The effect of parameter S choice on the on the accuracy of DMPA to discover known molecular associations in transcriptome data. The relative distance from the randomized median (A) and the corresponding P-value (B) of the validation score is shown.
**C-D:** The effect of parameter S choice on the runtime (C) of DMPA and amount of transcripts included in the complexes inferred with DMPA (D).
Mean ± SEM (n=3).

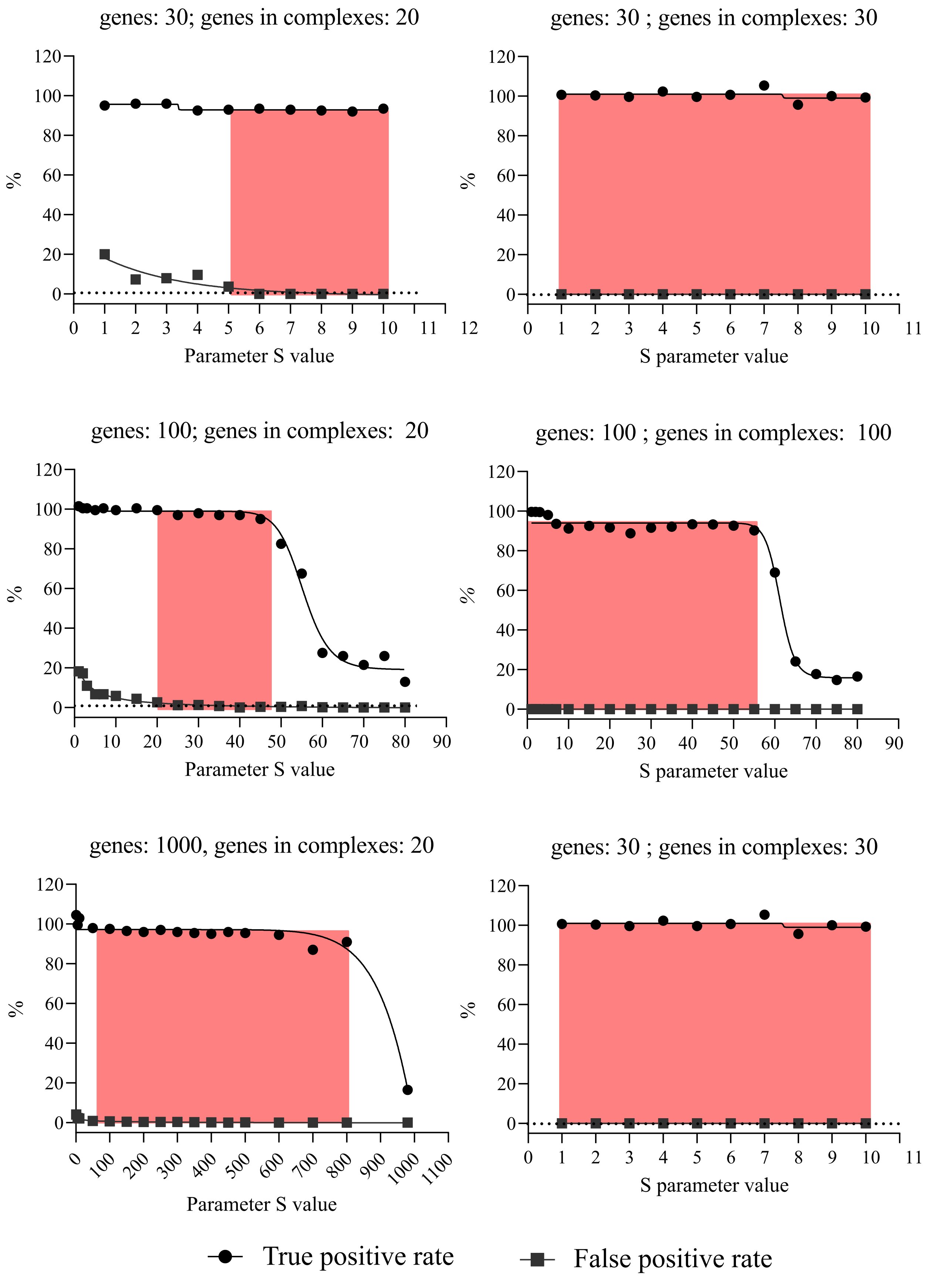


**Supplementary Figure 7. Effect of parameter S choice on the performance of DMPA with normally distributed simulated data.** The true and false postive rates at different parameter S values in datasets of different sizes and varying number of genes in true complexes. The optimal value range for the parameter S is indicated with red.

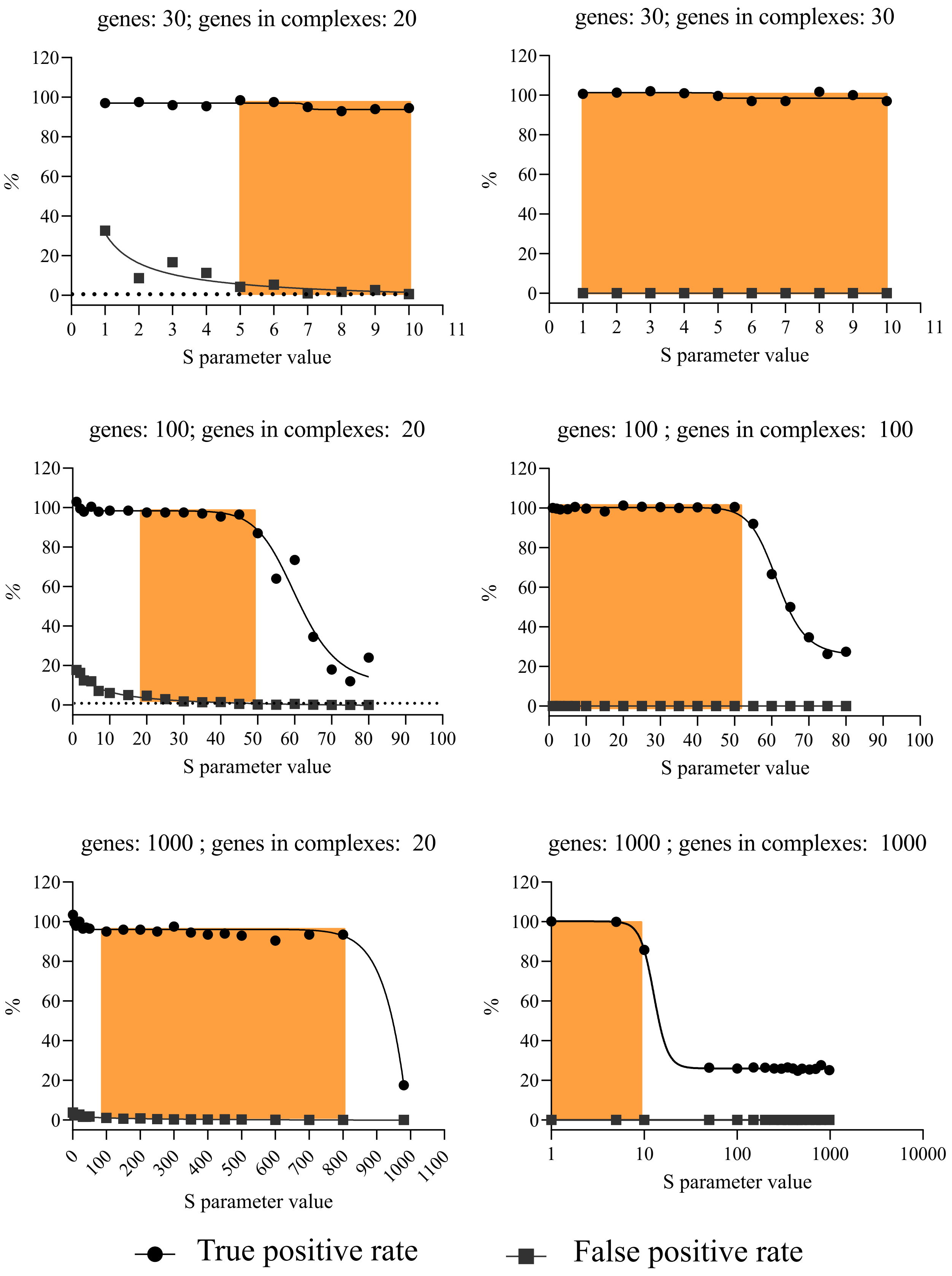


**Supplementary Figure 8. Effect of parameter S choice on the performance of DMPA with negative binomially distributed simulated data.** The true and false postive rates at different parameter S values in datasets of different sizes and varying number of genes in true complexes. The optimal value range for the parameter S is indicated with orange.

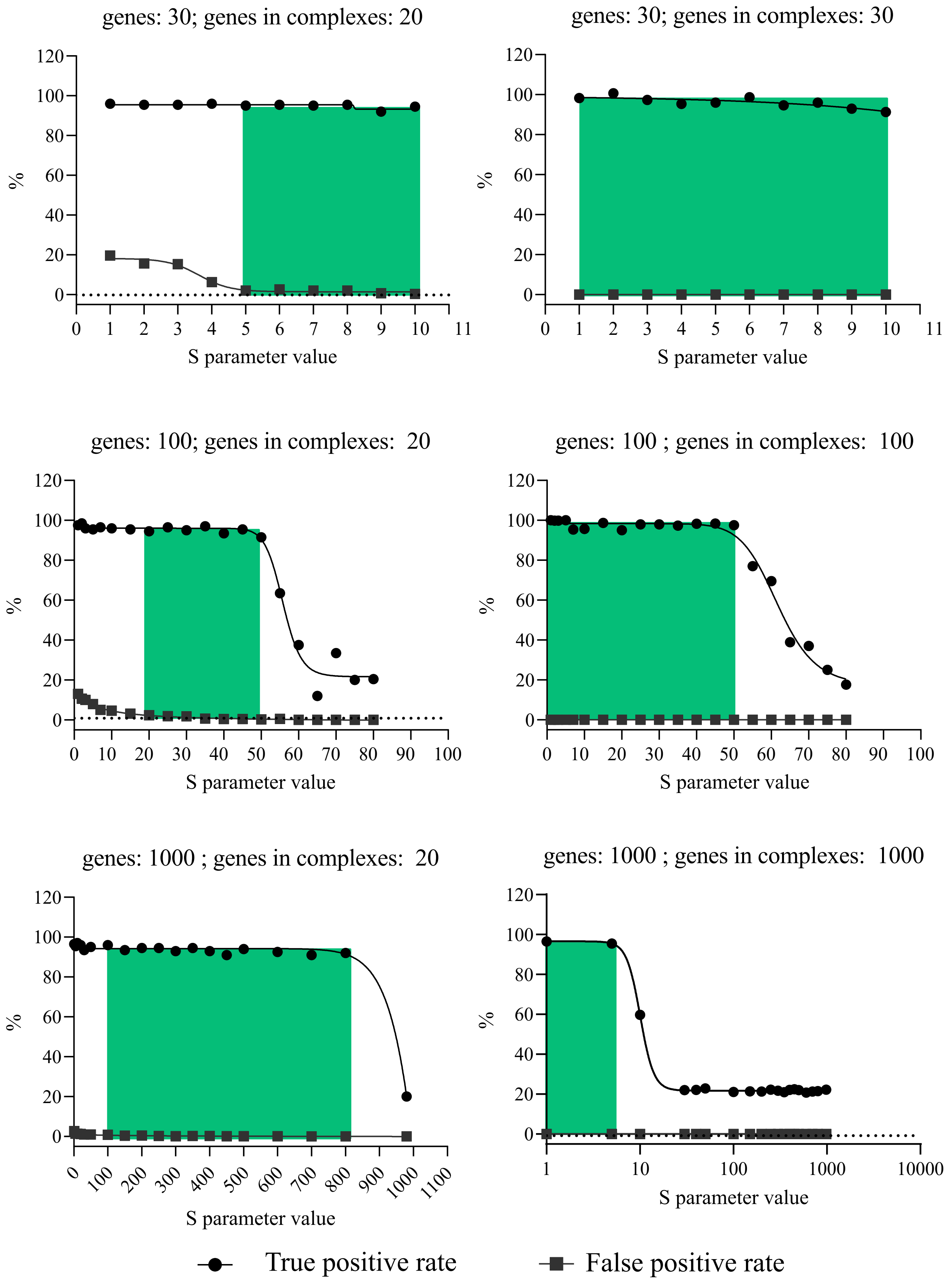


**Supplementary Figure 9. Effect of parameter S choice on the performance of DMPA with beta distributed simulated data.** The true and false postive rates at different parameter S values in datasets of different sizes and varying number of genes in true complexes. The optimal value range for the parameter S is indicated with turquiose.

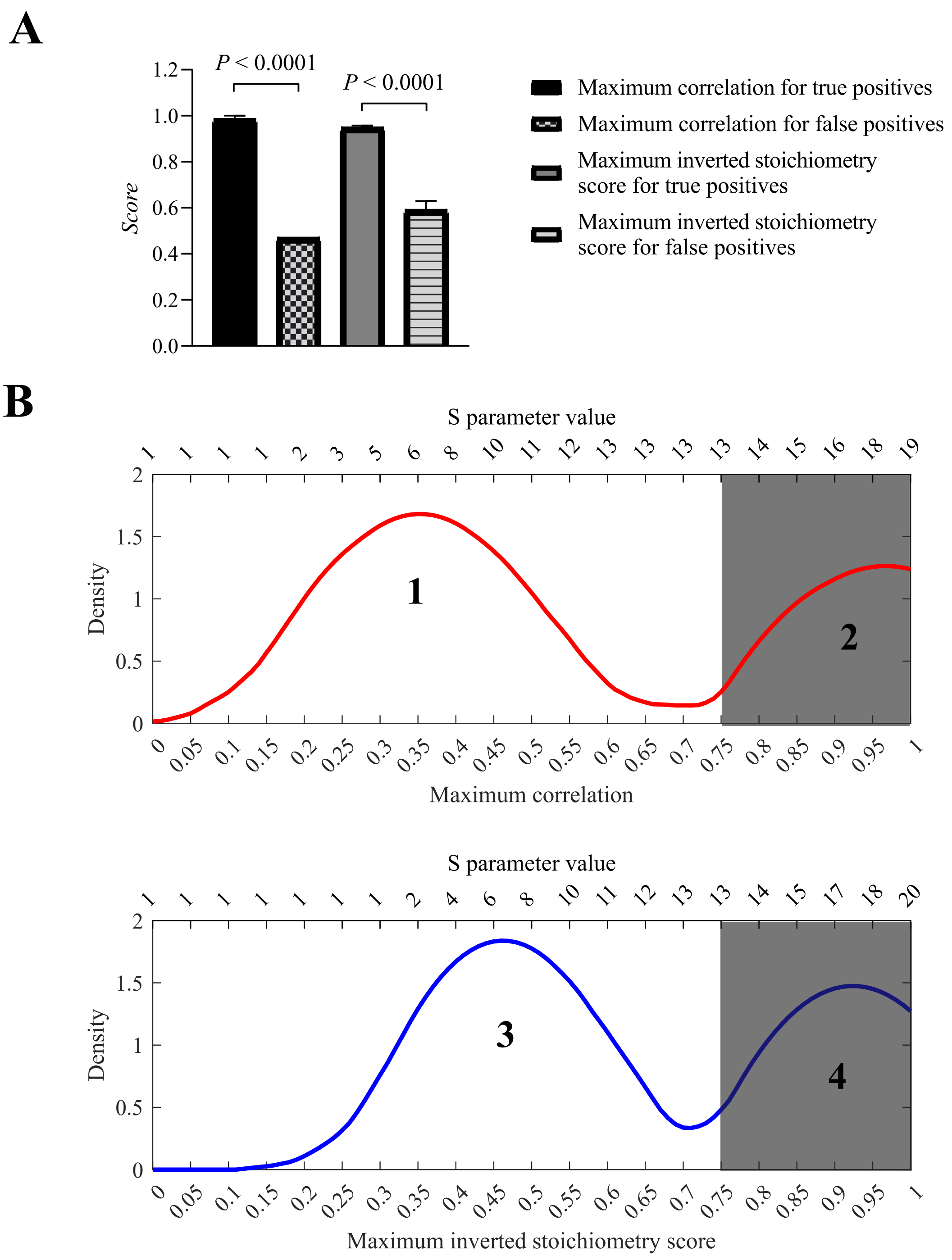


**Supplementary Figure 10. The determinants of accurate S parameter value choice.**
**A:** The median maximum correlation and inverted stoichiometry score for false and true positives in simulated data. Mean ± SD (n=30). Two-tailed Mann-Whitney U-test.
**B:** The output of the script designed to inform the S parameter value choice. Peak 1 and 3 represent the peaks for the false positives and peaks 2 and 4 the true positives in maximum correlation and inverted stoichiometry score, respectively. The grey area represent the simulated genes included in the analysis after cut-off at optimal parameter S value 13 is set for the algorithm for the simulated data. 20 true positives and 25 false positives were simulated.

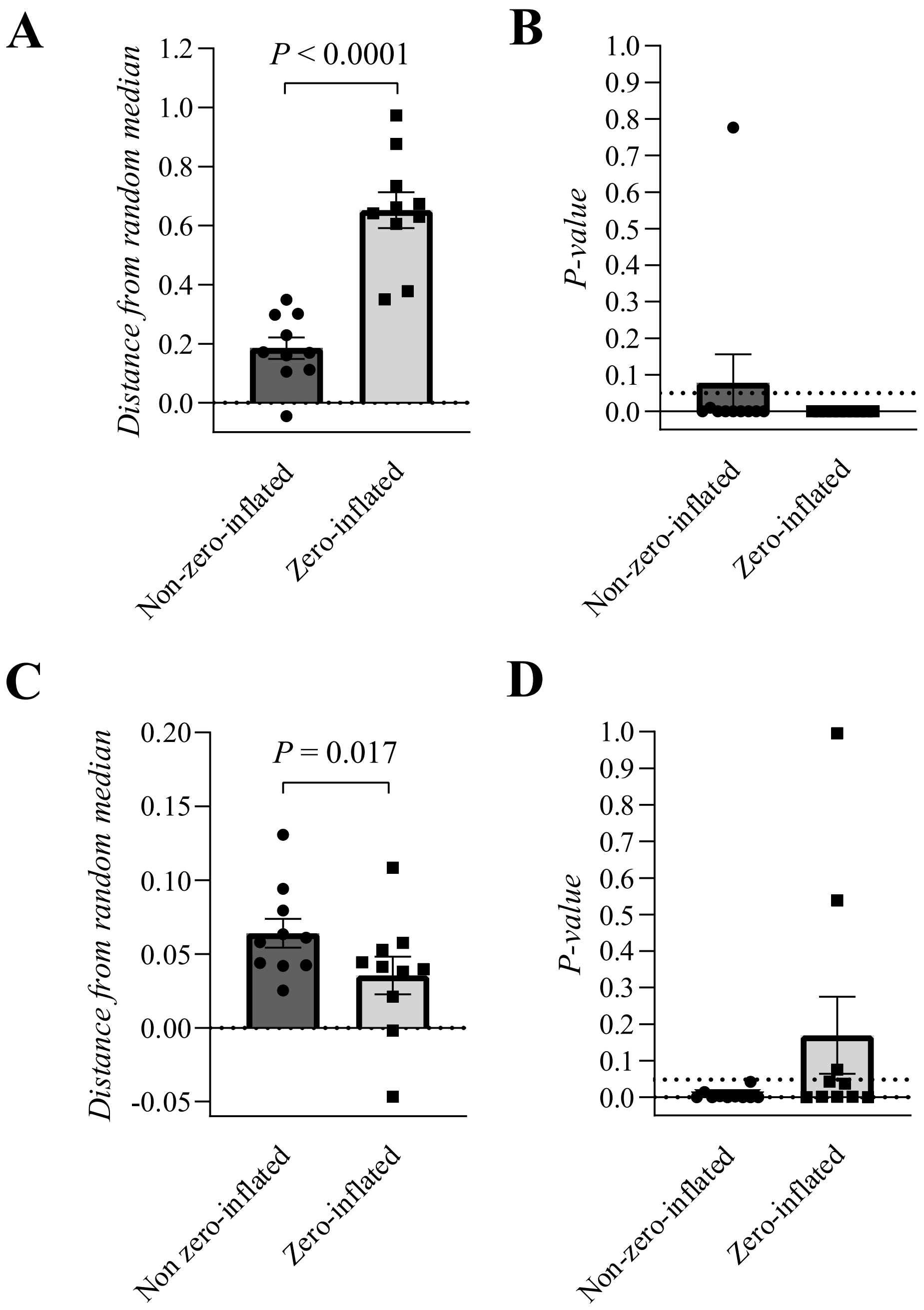


**Supplementary Figure 11. The effect of the zero-inflated version of DMPA on the performance of DMPA with zero-inflated and non-zero-inflated data.**

**A-B:** The effect of the zero-inflated version on the performance of DMPA to discover known protein-protein interactions in zero-inflated interactome data. The relative distance of the validation score from the randomized median and the corresponding P-value is shown.

**C-D:** The effect of the zero-inflated version on the performance of DMPA to discover known transcription factor target gene relationships in non-zero-inflated transcriptome data. The relative distance of the validation score from the randomized median and the corresponding P-value is shown.

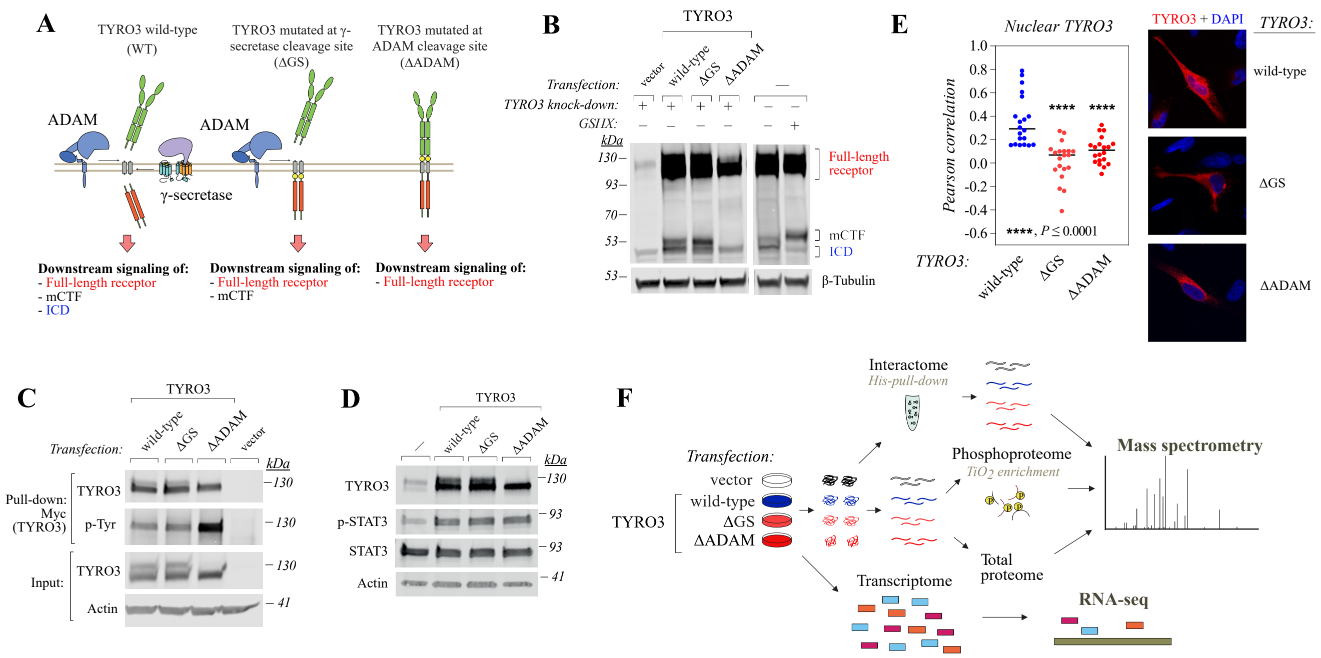


**Supplementary Figure 12. Functional validation of cleavage-resistant TYRO3 receptor variants.**
**A:** A Schematic depicting the signaling mediated by wild-type and regulated intramembrane proteolysis (RIP)-resistant variants (ΔGS, ΔADAM) of TYRO3. Since the cleavage events in RIP are sequential, blocking only the gamma-secretase cleavage will allow the receptor still to signal through the membrane-anchored C-terminal fragment (mCTF) in addition to the full-length receptor. Both cleavage events are blocked when the primary shedding performed by ADAM proteases is inhibited leading the receptor to signal only through the canonical full-length receptor. The wild-type receptor in turn is able to signal through the canonical full-length receptor, the mCTF and the released soluble intracellular domain (ICD).
**B:** Western analysis of WM-266-4 cells expressing the indicated TYRO3 variants and treated with or without the gamma-secretase inhibitor GSI IX. The full-length receptor and the different cleavage products of TYRO3 are indicated.
**C:** Western analysis of the autophosphorylation of TYRO3 in WM-266-4 cells expressing the indicated TYRO3 variants.
**D:** Western analysis of the phosphorylation status of TYRO3 downstream effector STAT3 in WM-266-4 transfectants.
**E:** Confocal microscopy analysis of TYRO3 in WM-266-4 cells expressing the indicated V5-tagged variants of TYRO3. V5 signal is shown in red and DAPI-stained nuclei in blue. Nuclear localization is presented as Pearson correlation coefficient of TYRO3-V5 co-localizing with DAPI within the cells. For statistical testing, the non-parametric Kruskal-Wallis ANOVA was utilized. The post hoc analyses were conducted with the Mann-Whitney U test and the resulting P-values were corrected with the method of Benjamini, Krieger and Yekutieli. One dot represents one cell and the horizontal line the median value.
**F:** The workflow of interactome, phosphoproteome, proteome and transcriptome data acquisition from WM-266-4 transfectants.

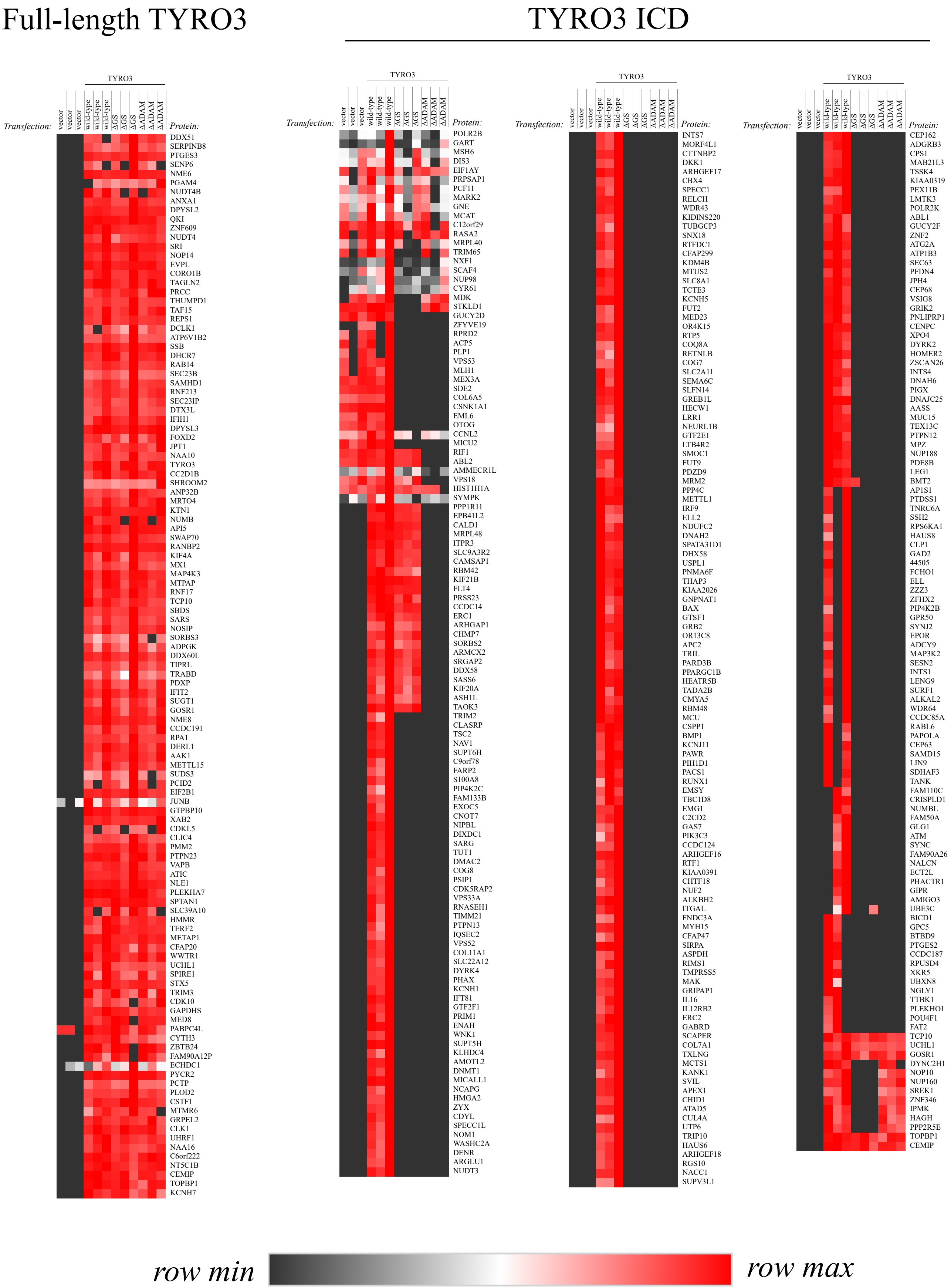


**Supplementary Figure 13. Interactome of full-length TYRO3 and cleaved TYRO3 ICD.** The TYRO3 precipitates of WM-266-4 transfectants were analyzed with mass spectrometry. Differential expression analysis was conducted to discover the proteins co-precipitating with the full-length TYRO3 and the cleaved TYRO3 ICD. Values are presented in a relative scale. For the original values, please see Supplementary Table 6. ICD: intracellular domain.

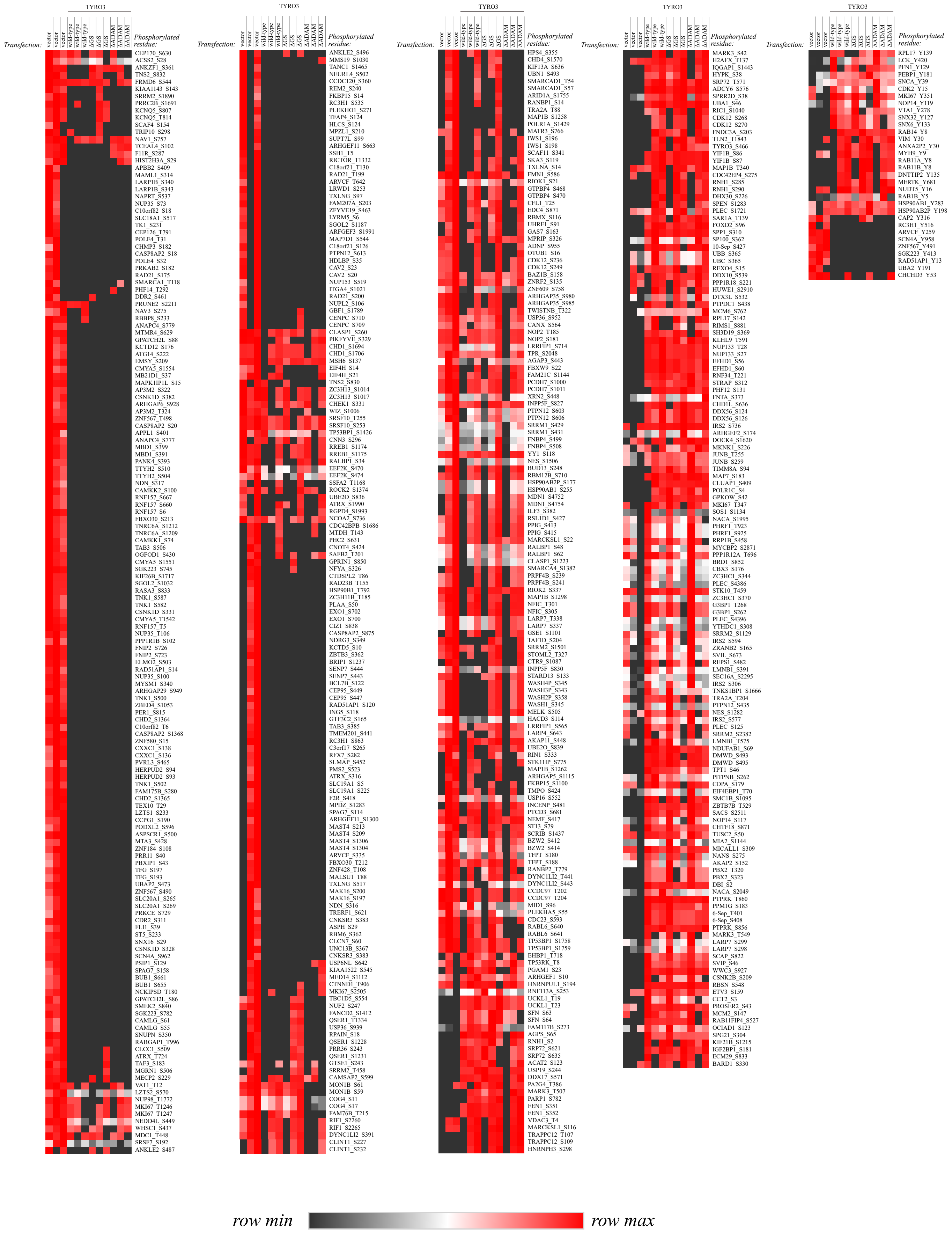


**Supplementary Figure 14. Differentially expressed phosphoproteome of full-length TYRO3.** The phosphoproteome of WM-266-4 transfectants was acquired with mass spectrometry. Differential expression analysis was conducted to discover the differentially phosphorylated residues associated with the signaling of full-length TYRO3. Values are presented in a relative scale. For the original values, please see Supplementary Table 7.

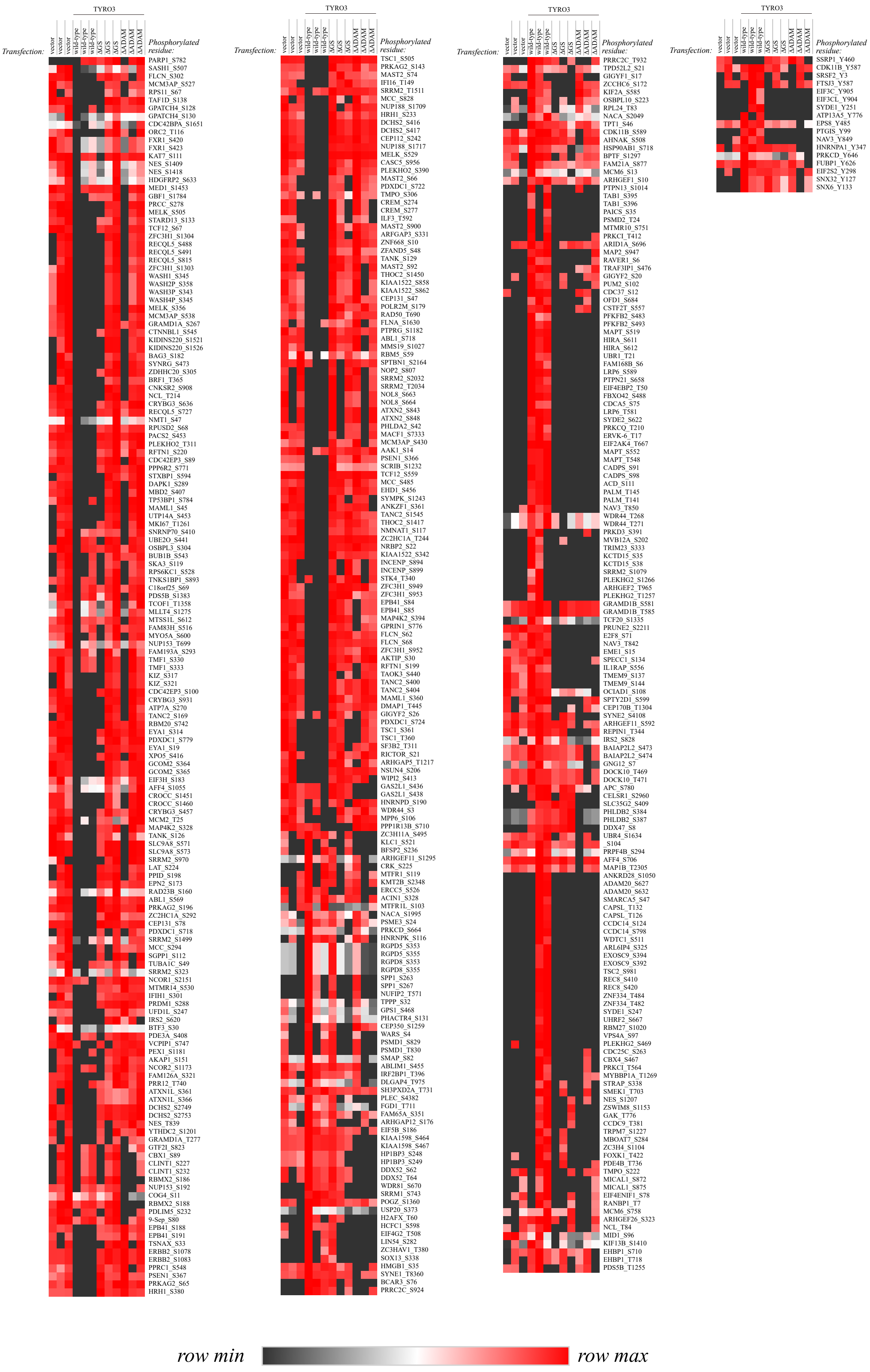


**Supplementary Figure 15. Differentially expressed phosphoproteome of cleaved TYRO3 ICD.** The phosphoproteome of WM-266-4 transfectants was acquired through mass spectrometry. Differential expression analysis was conducted to discover the differentially phosphorylated residues associated with the signaling of TYRO3 ICD. Values are presented in a relative scale. For the original values, please see Supplementary Table 7. ICD: intracellular domain.

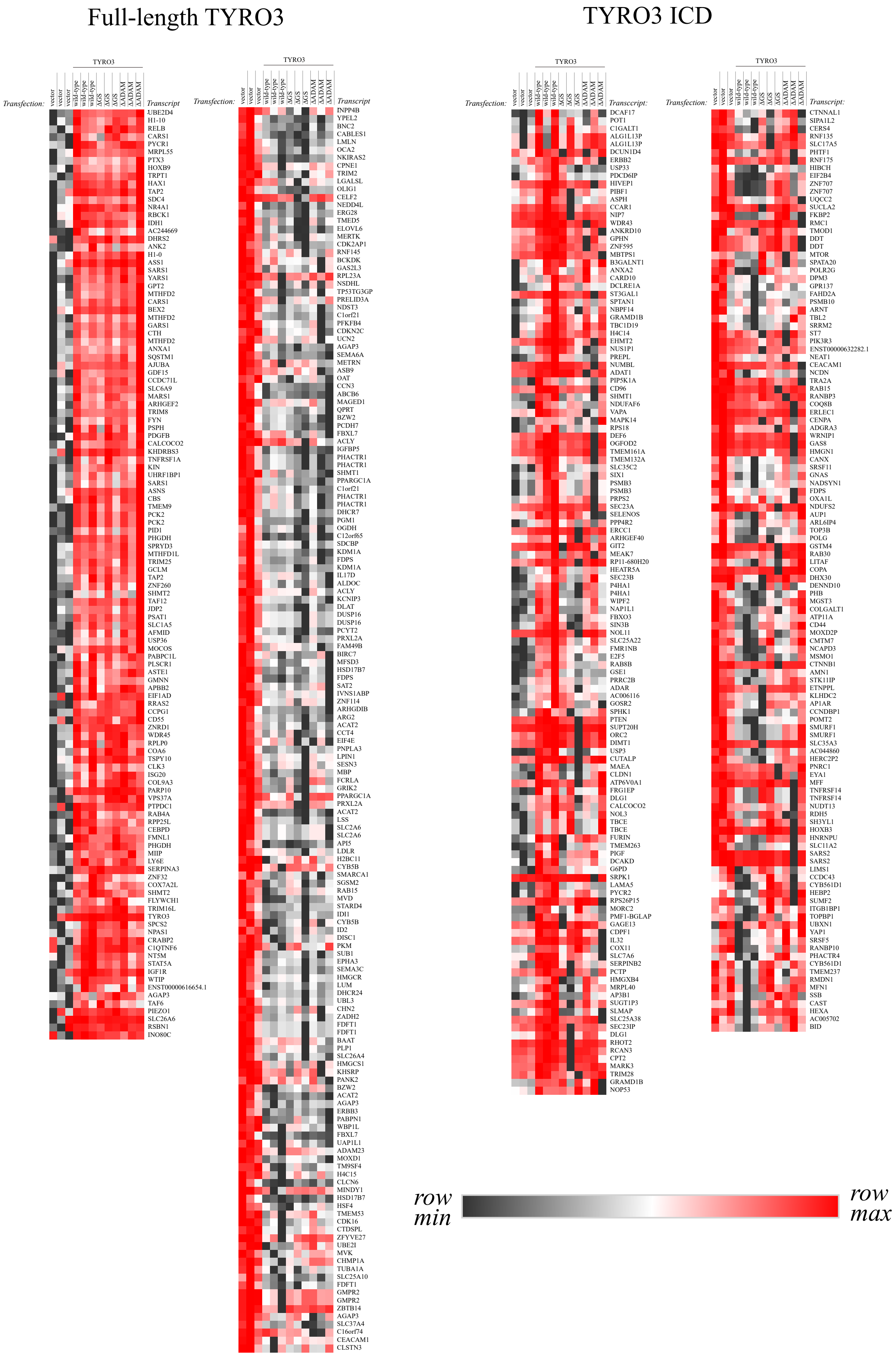


**Supplementary Figure 16. Differentially expressed transcriptome of full-length TYRO3 and cleaved TYRO3 ICD.** The mRNA extracts of WM-266-4 transfectants were sequenced with Illumina HiSeq. Differential expression analysis was conducted to discover the differentially expressed transcripts associating with full-length TYRO3 and the TYRO3 ICD. The values are presented in relative scale. For the original values, please see Supplementary Table 8. ICD: intracellular domain.

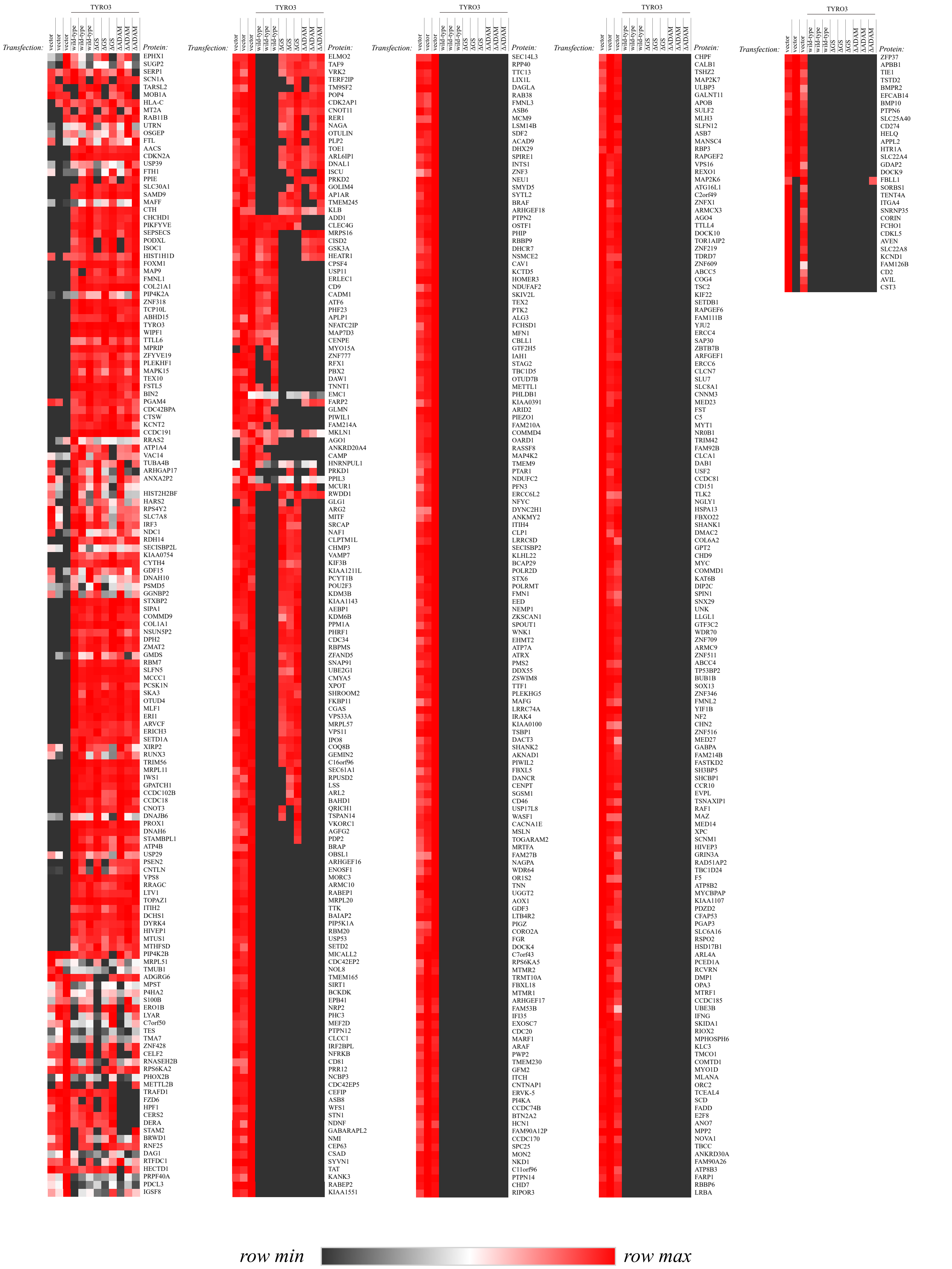


**Supplementary Figure 17. Differentially expressed proteome of full-length TYRO3.** The proteome of WM-266-4 transfectants was acquired with mass spectrometry. Differential expression analysis was conducted to discover the differentially expressed proteins associated with the signaling of full-length TYRO3. Values are presented in a relative scale. For the original values, please see Supplementary Table 9.

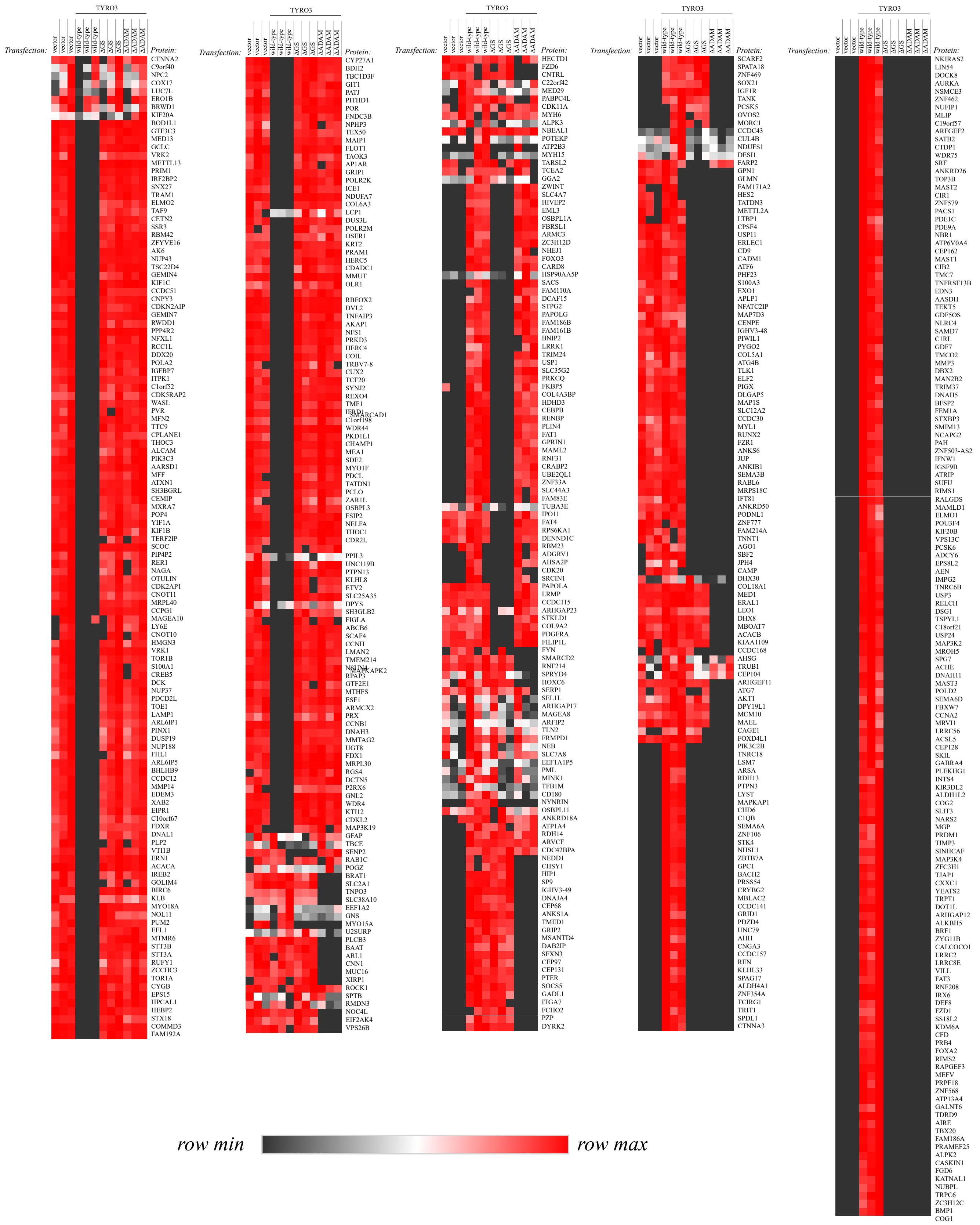


**Supplementary Figure 18. Differentially expressed proteome of cleaved TYRO3 ICD.** The proteome of WM-266-4 transfectants was acquired through mass spectrometry. Differential expression analysis was conducted to discover the differentially expressed proteins associated with the signaling of TYRO3 ICD. Values are presented in a relative scale. For the original values, please see Supplementary Table 9. ICD: intracellular domain.

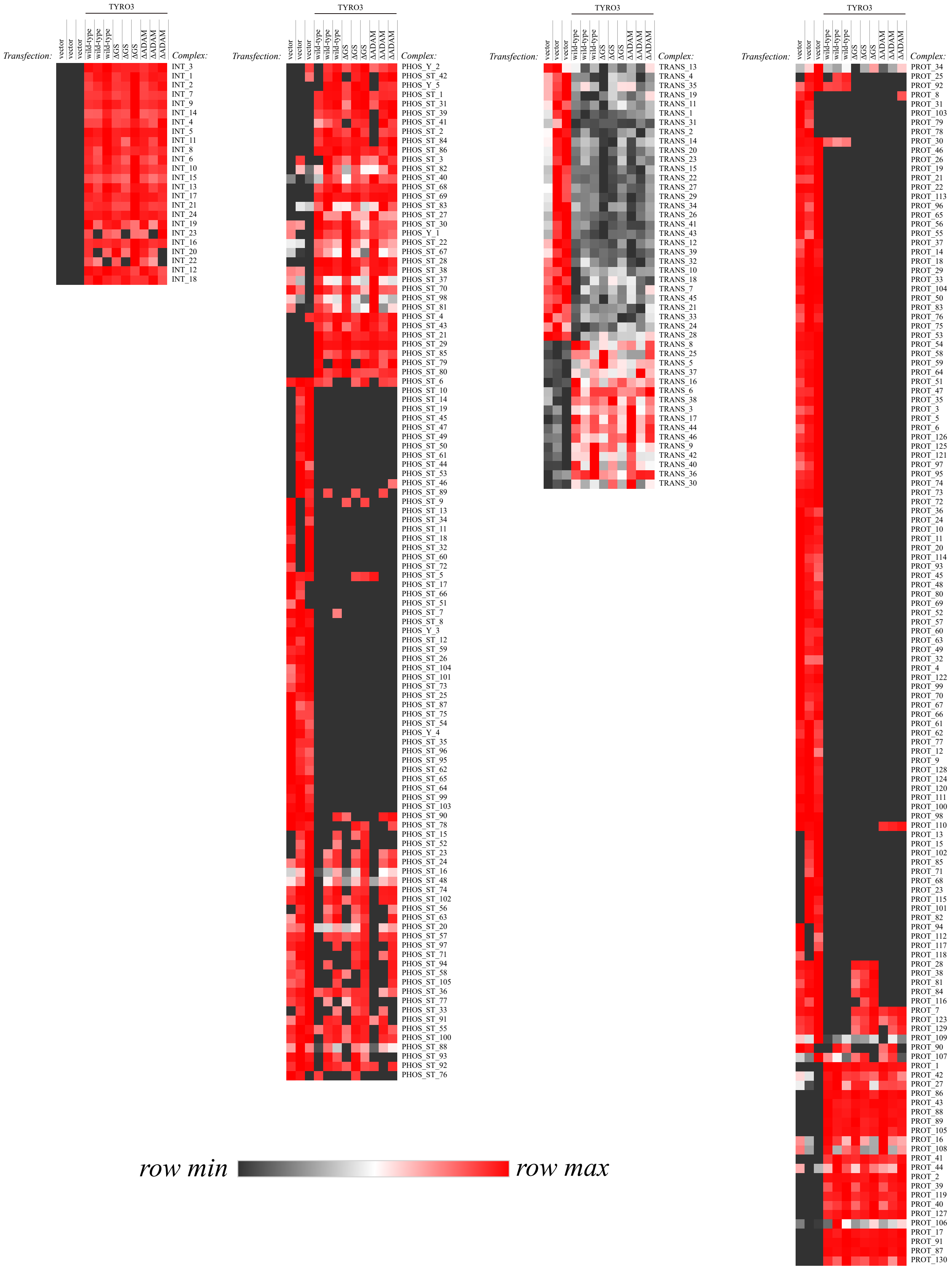


**Supplementary Figure 19. Regulatory complexes of full-length TYRO3.** The median expression of the regulatory complexes inferred with DMPA from the multi-omics data of full-length TYRO3-associated signaling in WM-266-4 transfectants. The presented values are in relative scale. For the original values, please see Supplementary Table 11.

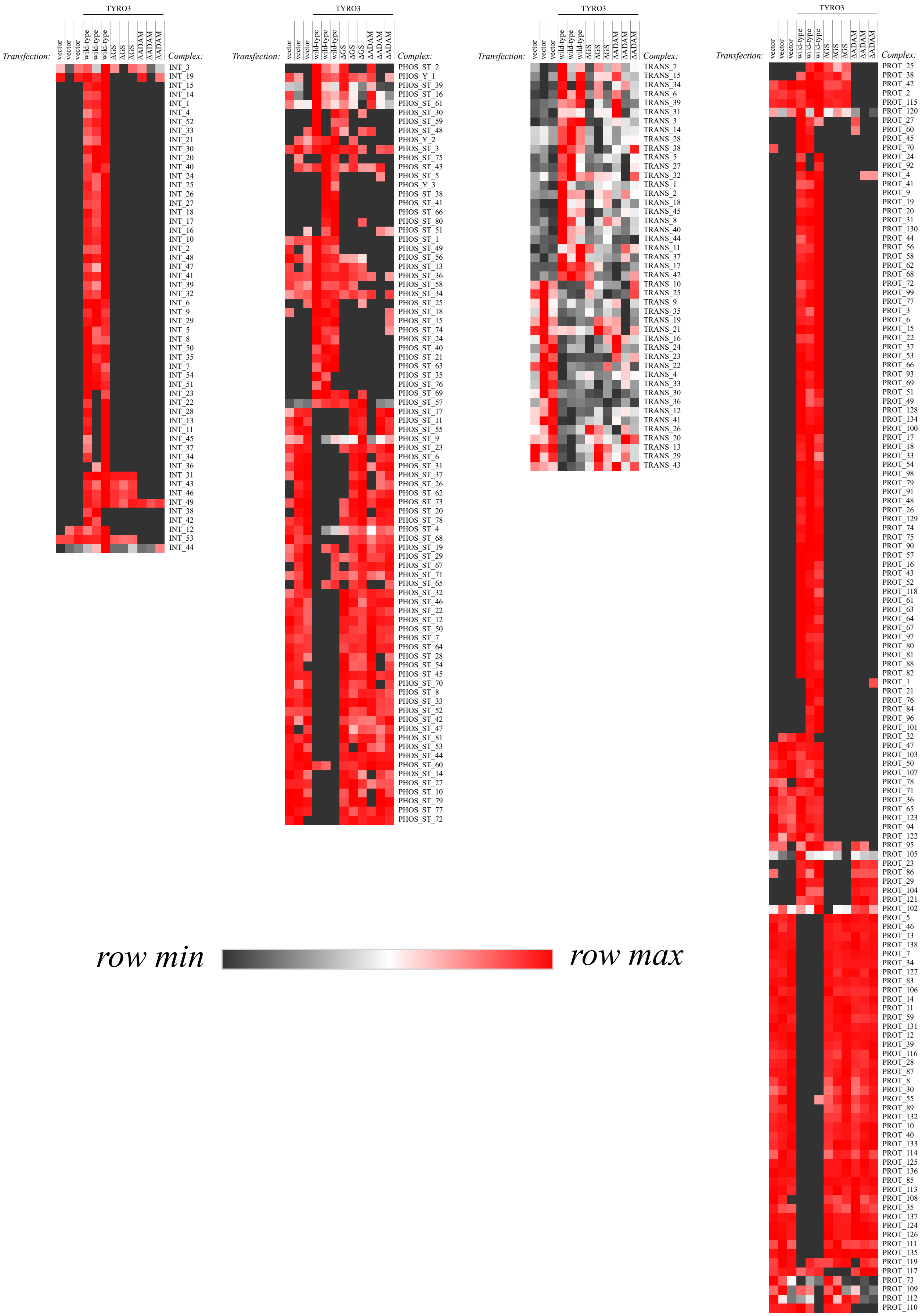


**Supplementary Figure 20. Regulatory complexes of cleaved TYRO3 ICD.** The median expression of the regulatory complexes inferred with DMPA from the multi-omics data of cleaved TYRO3 ICD-associated signaling in WM-266-4 transfectants. The presented values are in relative scale. For the original values, please see Supplementary Table 11. ICD: intracellular domain.

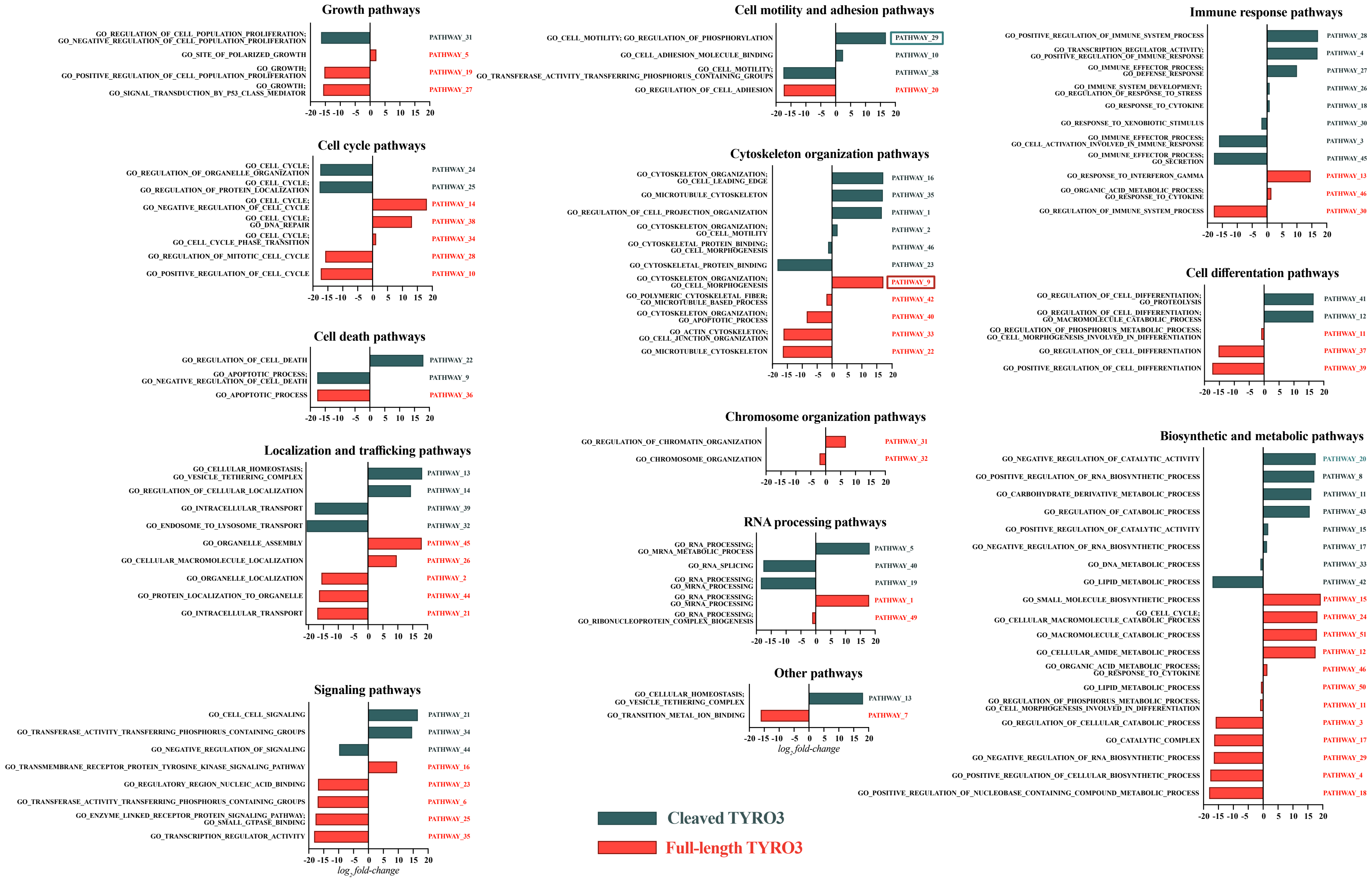


**Supplementary Figure 21. The functional categorization of the full-length and cleaved TYRO3 ICD pathways inferred with the DMPA.** The predicted function of the multi-omics pathways of the full-length and ICD of TYRO3 inferred with the de novo multi-omics pathway analysis. The bars represent the median pseudolog2 fold change of all the proteins, transcripts and phosphorylated residues in the pathway against the control condition. See supplementary tables 1 and 2 for further details on the pathways.

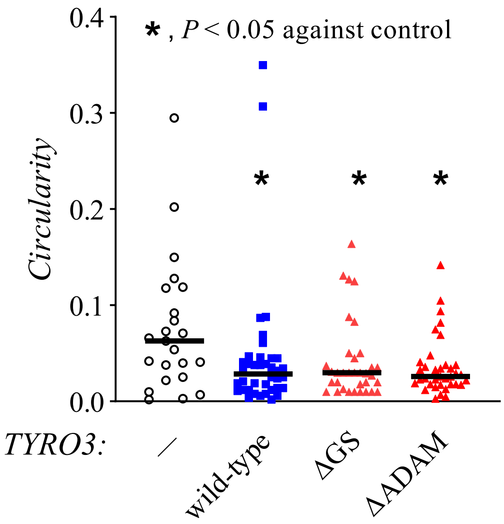


**Supplementary Figure 22**. **Morphology of WM-266-4 transfectants.** Two-dimensional morphological analysis of WM-266-4 transfectants. Circularity of the cells was measured from confocal images taken in plane with the plasma membrane with MorphoLibJ plugin of ImageJ. For statistical testing, the non-parametric Kruskal-Wallis ANOVA was utilized. The post hoc analyses were conducted with the Mann-Whitney U test and the resulting P-values were corrected with the method of Benjamini, Krieger and Yekutieli. One dot represents the morphology of one cell and the horizontal line the median value. ΔADAM, ADAM cleavage mutant; ΔGS, gamma-secretase cleavage mutant.
