## Supplementary Materials and Methods for "De Novo multi-omics pathway analysis (DMPA) designed for prior data independent inference of cell signaling pathways"

### Cell culture and transfections

WM-266-4 human melanoma cancer cells and HEK293T cells were cultured in DMEM medium (Life Technologies) supplemented with 10% (weight/vol) fetal calf serum (FCS) (Promocell). For the transfection of WM-266-4 cells, jetPRIME (Polyplus-transfection) transfection reagent was used according to manufacturer's protocol. For the production of lentiviral particles, the HEK293T cells were transfected with the lentiviral packaging and shRNA plasmids with FuGENE6 (Roche) according to the manufacturer's instructions.

### ADAM10 and ADAM17 cleavage site prediction

The cleavage site prediction method was modified from a previously published transcription factor binding prediction method (Lähdesmäki *et al*, 2008). The positional relative frequencies of the amino acid sequences in putative ADAM10 and ADAM17 cleavage sites were calculated from the results of two published peptide screens [2,3]. The cleavage sites for ADAM10 and ADAM17 were predicted with a sliding window analysis calculating the probability of a sequence window of 10 residues having an ADAM cleavage motif based on the positional relative frequencies of amino acid residues with a $0^{th}$ order Markov chain. The probability for lacking an ADAM cleavage motif was calculated from the same sequence windows with $0^{th}$ order Markov chain using the relative frequencies of amino acid residues in proteins as determined by Uniprot [4] as positional frequencies. The sequence windows with at least 2 times higher probability for containing an ADAM cleavage motif than lacking an ADAM cleavage motif were considered probable cleavage sites. Additional constraint for the cleavage motifs to reside within 35 amino acids from the transmembrane domain was added, due to known ectodomain size constraints of gamma-secretase substrates [5]. The method was validated by ensuring its capability in finding published cleavage sites in the peptides described in Goth *et al*, 2015. P408/L409 and A418/G419 of TYRO3 were predicted to serve as cleavage sites with highest predicted relative cleavage site probabilities for both ADAM10 and ADAM17.

### Plasmids and cloning

The pDONR223-TYRO3ΔGS vector including a TYRO3 insert with a mutated gamma-secretase cleavage site (I449A; TYRO3 ΔGS) has been previously described (Merilahti *et al*, 2017). Constructs encoding TYRO3 with mutated ADAM cleavage sites (P408A/L409P/G419P triple mutation; TYRO3 ΔADAM) were generated using synthetic DNA fragments (Integrated DNA Technologies) and assembled into pDONR223-based TYRO3 Gateway entry plasmid (Addgene kit #1000000014) [8,9] using NEBuilder HiFi DNA Assembly Master Mix (New England Biolabs). Wild-type TYRO3, TYRO3 ΔGS and TYRO3

ΔADAM constructs were cloned from pDONR223 plasmids into pEZY-myc-his (Addgene #18791) [10] and pMAX-DEST (Addgene #37631) [11] expression plasmids using Gateway Cloning Technology with LR Clonase II Enzyme Mix (Life Technologies) to allow expression of C-terminally Myc-his-tagged and V5-tagged proteins, respectively. Mutations were verified by sequencing.

For the production of lentiviral particles carrying shRNAs, the following plasmids were used: pLKO.1-puromycin plasmid (containing the shRNA) (Sigma-Aldrich), pRSV-Rev, pMDLg/pRRE and pMD2.G (a gift from Didier Trono; Addgene plasmids #12253, #12251 and #12259). Restriction digestion with EcoRI was used to verify the integrity of the commercial plasmids.

### Antibodies

Anti-TYRO3 (5585), anti-V5 (13202), anti-phospho-STAT3 (8862), and anti-STAT3 (9972) antibodies were purchased from Cell Signaling Technology. Anti-actin (MA1-744) antibodies were purchased from Thermo Fisher Scientific. Anti-β-tubulin (T7816) and anti-phosphotyrosine (4G10; 5-321) antibodies were purchased from Sigma-Aldrich.

### Immunoprecipitation and Western analysis

For immunoprecipitation experiments, WM-266-4 vector control cells or cells expressing Myc-tagged TYRO3 constructs were incubated in serum-free conditions overnight before lysis into lysis buffer (0.1% Triton X-100, 10 mM Tris-Cl pH 7.4, 150 mM NaCl including Pierce protease and phosphatase inhibitor mini tablet (Thermo Fisher Scientific)). The cell lysates were pre-cleared with 50 μl Pierce protein G magnetic beads (Thermo Fisher Scientific) at 4°C for 1 hour and subjected to affinity enrichment with Pierce anti-c-Myc magnetic beads (Thermo Fisher Scientific) at 4°C overnight. The beads were washed five times with 5×TBS-T buffer (125 mM Tris, 750 mM NaCl, 0.25% Tween-20) and once with ultrapure water. The proteins were denatured and eluted from the beads by incubation at 95°C for 10 minutes in SDS-PAGE loading buffer.

For Western analyses, cell lysates were prepared as above and run on SDS–PAGE gels. The separated proteins were transferred to nitrocellulose membranes, which were incubated with indicated primary antibodies and IRDye-conjugated secondary antibodies (LI-COR) to label the proteins of interest. The IR-signals were detected with the Odyssey CLx imaging system (LI-COR).

### Generation of stable TYRO3 knock-down cell lines

For stable downregulation of TYRO3, TYRO3-targeted lentiviral shRNA plasmid TRCN0000231528 (CCGGTTGGTATCTCAGGTCTGAATCCTCGAGGATTCAGACCTGAGATACCAATTTTTG) (MISSION, Sigma-

Aldrich) was used. The control lentiviral shRNA plasmid (Addgene plasmid #1864) was a kind gift from David Sabatini [12]. Third generation lentiviral packaging system (Addgene) was used in HEK293T cells to produce shRNA-carrying lentiviruses. Growth medium was changed every 24 hours post-transfection. Virus-containing medium was collected after 48 and 72 hours, filtered and tittered. The virus-containing medium was used to infect WM-266-4 cells at a multiplicity of infection of 2 in the presence of 8 µg/ml Polybrene (Sigma-Aldrich). Cells were maintained in the presence of 1 µg/ml puromycin (Sigma-Aldrich) to select the cells that stably express the lentiviral shRNA plasmids.

### Immunofluorescence and confocal microscopy

To detect ectopically expressed V5-tagged TYRO3 constructs, WM-266-4 transfectants were cultured on coverslips in serum-free conditions overnight. The cells were fixed with 4% paraformaldehyde, and permeabilized with 0.1% Triton X-100. Cells were stained with anti-V5 (13202; Cell Signaling Technologies) and AlexaFluor 555 goat anti-rabbit (Molecular Probes). After labeling the nuclei with 4',6-diamid-ino-2-phenylindole (DAPI; Sigma-Aldrich), the cells were mounted with Mowiol 40-88 (Sigma-Aldrich). Images were acquired with Zeiss LSM 880 confocal microscope (Zeiss). EzColocalization plugin [13] in ImageJ (version 1.53c) was used to analyze colocalization from confocal slices taken from the middle of the nucleus. The colocalization of V5-signals with DAPI was calculated using Pearson correlation coefficient [14] to measure the nuclear localization of TYRO3.

### RNA-sequencing

The WM-266-4 cell transfectants were starved in serum-free medium overnight and lysed. Total RNA was extracted using NucleoSpin RNA Plus kit (Macherey-Nagel).

The quality of the RNA samples was ensured using Advanced Analytical Fragment Analyzer (Agilent). The sequencing library was created with 300 ng of sample with TruSeq Stranded mRNA HT Kit (Illumina) and indexed with IDT for Illumina TruSeq RNA UD Indices according to manufacturer's protocol. Genome-wide strand-specific RNA-sequencing (RNA-seq) was performed at Turku Bioscience Centre sequencing core with Illumina HiSeq3000 using 75 bp paired-end reading.

### Mass spectrometry sample preparation

The WM-266-4 cell transfectants were lysed in mass spectrometry lysis buffer (6 M guanidine hydrochloride, 100 mM Tris-HCl pH 8.5, 5 mM Tris(2-carboxyethyl)phosphine (TCEP), 10 mM chloroacetamide) or affinity enrichment lysis buffer (70 mM octyl-β-D-glucopyranoside, 25 mM Tris-HCl pH 7.5, 150 mM NaCl, Pierce protease and phosphatase inhibitor mini tablet). The lysates were centrifuged at 20000 x g for 6 minutes and the supernatants were collected. The protein concentration of the lysates was measured with Bradford protein assay (Bio-Rad) before proceeding to affinity enrichment or to protein digestion to peptides.

### Affinity enrichment of mass spectrometry samples

The WM-266-4 cell transfectants were subjected to protein crosslinking with 2 mM DTBP for 10 minutes. The crosslinking reaction was quenched by incubation with 50 mM Tris-HCl pH 7.5 for 15 minutes before cell lysis. Equal amounts of WM-266-4 cell lysates were pre-cleared with Pierce protein G magnetic beads (Thermo Fisher Scientific) at 4 °C for 1 hour and subjected to affinity enrichment with HisPur Ni-NTA magnetic beads (Thermo Fisher Scientific) at 4 °C overnight. The beads were washed five times with 5×TBS-T buffer (125 mM Tris, 750 mM NaCl, 0.25% Tween-20) and once with ultrapure water. The proteins were denatured, alkylated and eluted from the beads at 95 °C for 10 minutes in elution buffer (6 M guanidine hydrochloride, 100 mM Tris-HCl pH 8.5, 10 mM Tris(2-carboxyethyl)phosphine (TCEP), 10 mM chloroacetamide).

### Protein digestion to peptides

Proteins enriched in affinity enrichment or purified cell lysates were digested with lys-C (enzyme/protein ratio 1:100) for 1 hour at 37 °C. The samples were diluted 1:10 with 50 mM $NH_4HCO_3$ and digested with trypsin (enzyme/protein ratio 1:100) at 37 °C overnight.

### Sample desalting

Sep-Pak tC18 96-well plate (Waters) was activated with 100% methanol and conditioned with 0.1% TFA in 80% acetonitrile (Thermo Fisher Scientific). Digested peptides were acidified to a pH 3 with trifluoroacetic acid (TFA) and desalted with the activated and conditioned Sep-Pak tC18 96-well plate. The peptides were washed with 0.1% TFA and eluted with 0.1% formic acid in 50% acetonitrile. Samples were dried in Hetovac vacuum centrifuge (Heto Lab Equipment) and stored dry in -20 °C until analysis with mass spectrometer.

### Phosphopeptide enrichment

Phosphopeptides were enriched from desalted and dried peptides using Pierce high-select $TiO_2$ phosphopeptide enrichment kit (Thermo Fisher Scientific) according the manufacturer's instructions.

Following elution, phosphopeptides were dried with Hetovac vacuum centrifuge and stored as dry in -20 °C until analysis with mass spectrometer.

### Mass spectrometry

Dried peptide samples were resuspended in 0.1% formic acid and sample concentrations were measured with Nanodrop 1000 (Thermo Fisher Scientific). Equal amounts of samples were analyzed on an Easy-nLC 1000 coupled to a Orbitrap Fusion Lumos instrument (Thermo Fisher Scientific) at Turku Bioscience Centre Proteomics Core Services. Peptides were loaded on in-house packed 100 μm × 2 cm precolumn packed with ReproSil-Pur 5 μm 200 Å C18-AQ beads (Dr. Maisch) using 0.1 % formic acid in water (buffer A) and separated by reverse phase chromatography on a 75 μm × 15 cm analytical column packed with ReproSil-Pur 5 μm 200 Å C18-AQ beads (Dr. Maisch). All separations were performed using a 60 minute gradient ranging from 8% buffer B (80% acetonitrile in 0.1% formic acid) to 21% in buffer B in 28 minutes and to 36% buffer B in 22 minutes and ramped to 100% buffer B in 5 minutes at flow rate of 300 nl/minute. The washout followed at 100% buffer B for 5 minutes.

All MS spectra were acquired on the orbitrap mass analyzer and stored in centroid mode. For data-dependent acquisition experiments, full MS scans were acquired from 300 to 1600 m/z at 120,000 resolution with fill target of 7E5 ions and maximum injection time of 50 ms. The most abundant ions on the full MS scan were selected for fragmentation using 1.6 m/z precursor isolation window and beam-type collisional-activation dissociation (HCD) with 30% normalized collision energy for a cycle time of 3 seconds. MS/MS spectra were collected at 15,000 resolution with fill target of 5E4 ions and maximum injection time of 100 ms. Fragmented precursors were dynamically excluded from selection for 35 seconds.

### Protein identification and quantification

MS/MS spectra were searched with Metamorpheus (version 0.0.304) [15] against human proteome containing known post-translational modifications (downloaded from Uniprot on 19.2.2019). The mass spectrometry files were calibrated, possible post-translational modifications were searched and peptides and proteins were identified and quantified using FlashLFQ algorithm [16]. The following parameters were used for the post-translational modifications search. Cysteine carbamidomethylation and methionine oxidation were set as constant and variable modifications, respectively and G-DPM search in Metamorpheus was additionally used to discover other modifications. Search results were filtered to a 1% FDR at PSM level. Peptides were accepted with search engine score above 5.

### RNAseq data analysis

The RNAseq reads were quality checked with FastQC (Babraham Bioinformatics), quality and adapter trimmed with PRINSEQ [17] and Trimmomatic [18], and pseudoaligned with kallisto v 0.46.0 [19] to human transcriptome Ensembl v96 [20] to retrieve TPM (transcripts per million) values. The batch effect between experiment 1 and experiments 2 and 3 was corrected with batchelor [21] and the TPM values were normalized by library size. Differential expression between the vector control sample and samples representing the different variants of TYRO3 was analyzed with DeSeq2 [22]. Transcripts with fold change over 1.5 and FDR-adjusted P-value lower or equal than 0.05 were chosen for further analysis. The transcripts significantly different between the vector control sample and all of the samples representing the TYRO3 variants (wild-type, ΔADAM or ΔGS) were considered to reflect the signaling mediated by the full-length TYRO3 receptor. The transcripts significantly different between the vector control sample and the sample representing wild-type TYRO3, but not between the vector control sample and the samples representing ΔADAM or ΔGS variant of TYRO3, were considered to reflect the signaling mediated by the soluble ICD of TYRO3.

### Mass spectrometry data analysis

The interactome, proteome and phosphoproteome data were normalized to the sum of intensities of all detected proteins in the sample. A probability density function was fitted with Epanechnikov kernel to median normalized intensities of different treatments to estimate the P-value for differential expression from the cumulative density function. Proteins with fold change over 1.5 and FDR adjusted P-value lower or equal to 0.05 were chosen for further analysis. The coprecipitating proteins, proteins and phosphorylated residues significantly different between the vector control sample and all of the samples representing the TYRO3 variants (wild-type, ΔADAM or ΔGS) were considered to reflect the signaling mediated by the full-length TYRO3 receptor. The proteins significantly different between the sample representing wild-type TYRO3 and the vector control sample as well as between the sample representing wild-type TYRO3 and samples representing the ΔADAM or ΔGS variant of TYRO3, were considered to reflect the signaling mediated by the soluble ICD of TYRO3.

### Data simulation

Simulation strategy adapted from Zuo et al. (2014) was utilized to simulate modules [23]. The relationships of complexes $X1=\{x1, x2, x3, x4, x5\}$ and $X2=\{x6, x7, x8, x9, x10\}$ were modeled as: $x1=s1+e1$, $x2=\lambda1 \cdot x1+e2$, $x3=\alpha1 \cdot x2+e3$, $x4=\alpha2 \cdot x3+e4$, $x5=\alpha3 \cdot x4+\lambda2 \cdot x1+e5$, $x6=\mu1 \cdot x10+e6$, $x7=\alpha4 \cdot x6+e6$, $x8=\alpha5 \cdot x7+e8$, $x9=\alpha6 \cdot x8+\mu2 \cdot x10+e9$, $x10=s2+e10$. In the normally distributed simulation data $s1,s2 \sim N(Y1,1)$ , $e1\text{-}e10 \sim N(1,Y1/10)$, where $Y1 \sim Z \in [2,30]$ and $\lambda1,\lambda2,\alpha1\text{-}\alpha6,\mu1,\mu2$ were set to 1. In the negative binomial simulation data $s1, s2 \sim NB(Y2,0.05)$, $e1\text{-}e10 \sim NB(Y2,0.5)$, where $Y2 \sim Z \in [2,15]$ and $\lambda1,\lambda2,\alpha1\text{-}\alpha6,\mu1,\mu2$ were set to 1. In the beta simulation data $s1, s2 \sim B(Y3,2)$, $e1\text{-}e10 \sim B(Y3,50)$, where

Y3 ~ $Z \in [2,10]$ and λ1,λ2,α1-α6,μ1,μ2 were set to 1. The datapoints with no true complex assignment were modelled as s3 – e11, where s3 ~ $Z \in [1,35]$, e11~N(0,1) or s3 ~NB(Y4,0.05), e11 ~NB(Y4,0.5), Y4 ~ $Z \in [2,15]$ or s3 ~B(Y4,3), e11~B(Y4,50), Y4 ~ $Z \in [2,10]$, respectively. 50 samples were simulated.

### Transcription factor and kinase prediction

An enrichment analysis to predict a transcription factor for each transcriptome and a kinase for each phosproteome regulatory complex was devised. Transcription factor target gene annotations from the ChEA3 data resource [24] were used and only the target gene annotations of the transcription factors present in the dataset with at least 5 transcripts were considered. The annotations of the upstream kinase for each phosphosite was acquired and combined from the results of a kinase knock-down screen [25], RegPhos 2.0 [26], PhosphoSitePlus [27], and KinaseNET [28] databases. The annotations of upstream kinases based on protein-protein interactions were accumulated from BioGrid [29], Cheng et al. [30], Harmonizome [31], HIPPIE [32], Mentha [33], Mint [34], PIPs [35], PSOPIA [36], Reactome [37], and STRING [38] databases. The kinase annotations were filtered according to the LFQ intensity of the kinases in the proteome and phosphoproteome data. Only substrate phosphosite or protein-protein interaction annotations of the kinases that were present in the corresponding samples with at least the LFQ intensity of 10000 were considered.

The enrichment analyses were created to follow the 3 steps as described: Step 1) The annotation most enriched in all the transcripts or phosphosites in all the complexes is given to the complexes for which the annotation gives the highest statistically significant enrichment score (the number of transcripts or phosphorylated residues found in the annotation divided by the number of transcripts or phosphorylated residues in the complex; P-value equal or less than the set threshold). The most enriched annotation and the complexes for which the annotation gives the highest enrichment score are removed. For the remaining dataset the same process is repeated until no complex remains. If more than one annotation provides the same overall and complex specific enrichment, all annotations are given as equally likely to the complexes. To derive a P-value for the enrichment of each annotation in each regulatory complex 10,000 random sets of equal size from all transcripts or phosphorylated residues identified in the transcriptome or phosphoproteome data are drawn. The respective enrichment of each of these randomized sets is derived and a probability density function is fitted with an Epanechnikov kernel to the enrichment scores. The P-values for the enrichment are drawn from the corresponding empirical cumulative distribution function. Step 2) Secondary transcription factor and kinase predictions are performed by assessing whether any of the primary transcription factors or kinases would be statistically significantly enriched also in the regulatory complexes were another transcription factor or kinase is assigned as the primary. P-values are similarly acquired for the secondary transcription factors and kinases. If more than one annotation provides the same complex specific enrichment with a P-value lower than 0.05, all annotations are given as equally likely to the complex. Step 3) If any unannotated complexes remain, all remaining annotations are searched to find the first statistically significant annotation with the highest enrichment score. If more than one annotation provides the same enrichment score with a P-value lower than the set threshold, all annotations are given as equally likely to the complex.

### Subcellular location prediction

An enrichment analysis to predict the subcellular location of each regulatory interactome complex was devised. The annotations of subcellular locations was acquired from the knowledge, experiments and text mining channels of COMPARTMENTS database [39]. The enrichment score (i.e., relative frequency) of each annotation for each interactome complex was calculated. To derive a P-value for the enrichment of each annotation in each regulatory complex, 1,000 random sets of equal size from all possible proteins identified in the interactome data were drawn. The respective enrichment of each of these randomized sets was derived and a probability density function was fitted with an Epanechnikov kernel to the enrichment scores. The P-values for the enrichment were drawn from the corresponding empirical cumulative distribution function. Annotations with a P-value lower or equal to 0.05 were considered significant. The subcellular location annotation with a highest enrichment score and lowest P-value were selected.

### Function prediction

An enrichment analysis to predict the biological process involved with the inferred pathways was devised. The Gene Ontology resource [40,41] annotations for biological processes were acquired from the MSigDB v7.3 [42]. The enrichment score (i.e., relative frequency) of each annotation for each pathway was calculated. To derive a P-value for the enrichment of each annotation in each pathway 10,000 random sets of equal size from all possible proteins, transcripts and phosphorylated proteins identified as significantly altered in the condition were drawn. The respective enrichment of each of these randomized sets was derived and a probability density function was fitted with an Epanechnikov kernel to the enrichment scores. The P-values for the enrichment were drawn from the corresponding empirical cumulative distribution function. Annotations with a P-value lower or equal to 0.05 were considered significant. The function annotation with a highest enrichment score and lowest P-value were selected. Contextually irrelevant annotations, such as specific functions of certain non-skin tissues or non-melanoma cell types, were ignored.


1. Lähdesmäki H, Rust AG, Shmulevich I. Probabilistic inference of transcription factor binding from multiple data sources. PLoS One. 2008;3. doi:10.1371/JOURNAL.PONE.0001820

2. Caescu CI, Jeschke GR, Turk BE. Active-site determinants of substrate recognition by the metalloproteinases TACE and ADAM10. Biochem J. 2009;424: 79–88. doi:10.1042/BJ20090549

3. Tucher J, Linke D, Koudelka T, Cassidy L, Tredup C, Wichert R, et al. LC-MS based cleavage site

profiling of the proteases ADAM10 and ADAM17 using proteome-derived peptide libraries. J Proteome Res. 2014;13: 2205–2214. doi:10.1021/pr401135u
